# A Pharmacophore Model for SARS-CoV-2 3CLpro Small Molecule Inhibitors and in Vitro Experimental Validation of Computationally Screened Inhibitors

**DOI:** 10.1101/2021.03.02.433618

**Authors:** Enrico Glaab, Ganesh Babu Manoharan, Daniel Abankwa

**Affiliations:** Luxembourg Centre for Systems Biomedicine (LCSB), University of Luxembourg, 7 avenue des Hauts Fourneaux, L-4362 Esch-sur-Alzette, Luxembourg; Department of Life Sciences and Medicine, University of Luxembourg, 7 avenue des Hauts Fourneaux, L-4362 Esch-sur-Alzette, Luxembourg

**Keywords:** COVID-19, SARS-CoV-2, pharmacophore, drug repurposing, 3CLpro, ligand activity assay, virtual screening, molecular dynamics simulation, machine learning

## Abstract

Among the biomedical efforts in response to the current coronavirus (COVID-19) pandemic, pharmacological strategies to reduce viral load in patients with severe forms of the disease are being studied intensively. One of the main drug target proteins proposed so far is the SARS-CoV-2 viral protease 3CLpro (also called Mpro), an essential component for viral replication. Ongoing ligand- and receptor-based computational screening efforts would be facilitated by an improved understanding of the electrostatic, hydrophobic and steric features that characterize small molecule inhibitors binding stably to 3CLpro, as well as by an extended collection of known binders.

Here, we present combined virtual screening, molecular dynamics simulation, machine learning and *in vitro* experimental validation analyses which have led to the identification of small molecule inhibitors of 3CLpro with micromolar activity, and to a pharmacophore model that describes functional chemical groups associated with the molecular recognition of ligands by the 3CLpro binding pocket. Experimentally validated inhibitors using a ligand activity assay include natural compounds with available prior knowledge on safety and bioavailability properties, such as the natural compound rottlerin (IC_50_ = 37 µM), and synthetic compounds previously not characterized (e.g. compound CID 46897844, IC_50_ = 31 µM). In combination with the developed pharmacophore model, these and other confirmed 3CLpro inhibitors may provide a basis for further similarity-based screening in independent compound databases and structural design optimization efforts, to identify 3CLpro ligands with improved potency and selectivity.

Overall, this study suggests that the integration of virtual screening, molecular dynamics simulations and machine learning can facilitate 3CLpro-targeted small molecule screening investigations. Different receptor-, ligand- and machine learning-based screening strategies provided complementary information, helping to increase the number and diversity of identified active compounds. Finally, the resulting pharmacophore model and experimentally validated small molecule inhibitors for 3CLpro provide resources to support follow-up computational screening efforts for this drug target.

## INTRODUCTION

The coronavirus disease 2019 (COVID-19), caused by the severe acute respiratory syndrome coronavirus 2 (SARS-CoV-2), represents a major challenge for health care systems around the world. Patients experience highly heterogeneous symptoms, ranging from asymptomatic forms and mild symptoms of a respiratory infection to severe illness, which can lead to hospitalization and death.

While the first vaccines against COVID-19 have received Emergency Use Authorization (EUA) by the Food and Drug Administration (FDA) and mass vaccination campaigns are currently ongoing, the available drug-based treatments for the disease are still limited. In the RECOVERY trial, a randomized trial designed to provide a fast and robust assessment of potential treatments for COVID-19, the corticosteroid dexamethasone was associated with a reduced mortality for patients with severe forms of the disease, when using a moderate dose (6 mg daily for 10 days) [1]. The FDA has included dexamethasone in their list of drugs used for hospitalized patients with COVID-19, and the European Medicines Agency (EMA) has endorsed the use of dexamethasone in patients from twelve years of age and weighing at least 40 kg, who require supplemental oxygen therapy. Moreover, the drug remdesivir, which inhibits the viral RNA-dependent RNA-polymerase (RdRp), was approved by the FDA for the treatment of COVID-19 requiring hospitalization. Remdesivir provided a statistically significant shorter median time to recovery in the clinical trial ACTT-1 [2], and significant symptom improvements vs. the standard of care in the GS-US-540-5774 trial [3], but no statistically significant improvements were observed for overall 29-day mortality in the ACTT-1 trial. In the independent SOLIDARITY trial no significant benefits were found in terms of reduced mortality, reduced initiation of ventilation or hospitalization duration [4]. Other potential treatment options against COVID-19 studied in clinical trials, such as tocilizumab [5], colchicine [6], synthetic neutralizing antibodies [7], or convalescent plasma [8], among others, were at the time of writing still under investigation. Overall, while first significant successes in COVID-19 treatment have been achieved using pharmacological methods, there is a broad consensus that additional primary or adjuvant treatment approaches are required to further reduce mortality and hospitalization rates.

For the development of new drug therapies against COVID-19 both human and viral protein drug targets have been proposed. Suggested human drug targets include in particular proteins involved in mediating the entry of SARS-CoV-2 into the cell, such as the proteases TMPRSS2 and FURIN, which have been implicated in the proteolytic processing of the viral spike protein before cell entry [9, 10]. The viral targets mainly include proteins involved viral replication (3CLpro / Nsp5 [11], PLpro / Nsp3 [12], RdRp / Nsp12 [13], Hel / Nsp13 [14]), and proteins associated with viral cell entry (S1, S2 [15]) and release (ORF3a [16, 17]).

One of the viral targets of particular interest is the 3C-like protease (3CLPro, Mpro), which is needed by SARS-CoV-2 for the cleavage of viral polyproteins into 11 non-structural proteins (NSPs) that are essential for viral replication [11]. Apart from its critical role in replication, 3CLPro may also represent a well-suited target for drug development, because multiple crystal structures are available with high resolution (< 1.3 Å) and favorable quality characteristics, and first small molecule inhibitors for both the SARS-CoV-2 and SARS-CoV forms of the protein have already been identified [18–21]. While these inhibitors still have limitations in terms of either their binding affinity for the target protein, their bioavailability, adverse drug effects or high manufacturing costs, the existing structural data for 3CLPro provides ample opportunities to apply both receptor- and ligand-based computational screening approaches in order to identify more potent and selective inhibitors.

To contribute to the ongoing research efforts on identifying improved 3CLPro inhibitors, here we present the results of a combined computational and experimental screening and pharmacophore generation approach. By integrating multiple computational approaches for molecular docking, ligand-based screening, machine learning and Molecular Dynamics (MD) simulations, we have ranked small molecule compounds from the public libraries provided by the repositories ZINC [22], SWEETLEAD [23] and the company MolPort (www.molport.com) as candidate inhibitors for 3CLPro. Top-ranked compounds and a few reference molecules previously proposed as 3CLpro inhibitors in the literature were experimentally assessed using a ligand activity assay to identify a subset of inhibitors and their IC_50_ values. We also compared the utility of different strategies for inhibitor discovery on the MolPort library. This extended analysis focused only on the MolPort compounds, because this library was our main resource for commercially available compounds and therefore of particular relevance for the experimental studies, and due to its smaller size (∼7.7 million compounds), an extension to more computationally demanding screening approaches was still feasible in terms of runtime requirements. Finally, the validated inhibitors, which include both natural compounds with existing information on ADMETox characteristics, as well as previously uncharacterized synthetic compounds, were used to generate a pharmacophore model (i.e. a computational model that describes the steric and chemical features of compounds associated with the binding to the target protein).

The information derived from this model and the confirmed inhibitors provide a resource to facilitate follow-up efforts on the structural design of more potent and selective 3CLpro inhibitors or the similarity-based inhibitor screening in additional compound databases.

The reader should note that the workflows used for computational screening in the present study are limited by their design for typical high-performance computing systems in an academic setting. Recent supercomputer-based virtual screening approaches, which have also been applied to 3CLpro inhibitor screening, can explore a significantly wider search space of compounds and conformations, and therefore also have the potential to achieve higher hit rates. Specifically, supercomputing-based screening methods have recently been shown to enable an improved modeling of 3CLpro receptor flexibility using ensemble docking [24], extensive MD simulations to generate high-resolution conformational ensembles [25], and the application of more advanced polarizable force fields [26]. Similarly, for the experimental assessment of 3CLpro inhibitory activity, recently developed assays published during the writing time of this manuscript provide a higher scalability and throughput than the Förster resonance energy transfer (FRET) assay used in the present work, and have already identified new 3CLpro inhibitors as drug candidates for further development [20].

Because supercomputing facilities and more recent inhibition assays were not available to us, the goal of our study was to contribute to research on inhibiting SARS-CoV-2 3CLpro by providing complementary results and data with the methods and hit rates achievable in an academic setting, in particular through the computational discovery and experimental confirmation of new micromolar 3CLpro inhibitors, and through the creation of pharmacophore models specific to the conformational and physicochemical properties of these inhibitors. The identified inhibitors and the information from the pharmacophore model provide both an input to support further screening studies, e.g. using independent private compound libraries, and a basis for the rational design of compounds with improved activity and ADMETox properties. The potential for further structural optimization of initial screening hits has recently already been illustrated in a study using computational lead optimization of the weak hit compound perampanel for 3CLpro inhibition, which led to the design and confirmation of multiple noncovalent, nonpeptidic 3CLpro inhibitors with significantly improved affinity in a kinetic assay, that were successfully confirmed to inhibit infectious SARS-CoV-2 replication *in vitro* [21, 27].

In the following sections, we first present the computational and experimental methods used for screening and compound validation, the results of the screening procedure in terms of the characteristics of the validated inhibitors, and finally a discussion of the pharmacophore model generated from the analysis of these confirmed inhibitors.

## METHODS

In order to identify and characterize small molecule inhibitors for SARS-CoV-2 3CLpro and build a pharmacophore model, a multi-step computational screening procedure was combined with subsequent ligand activity assay experiments. The main component of the computational screening consisted of a filtering procedure involving multiple receptor- and ligand-screening approaches in combination with a final molecular dynamics (MD) simulation to confirm the stability of the binding for the top-ranked compounds and docking poses. An overview of this part of the screening approach is shown in Figure 1, the detailed methodology, including the prior preprocessing of protein and compound structures, is discussed in the following sections below. Additionally, as part of the *in silico* selection of candidate compounds for experimental validation, the structural bioinformatics based screening was complemented by a machine learning approach for compound selection, using a training set of molecular descriptor data derived from known binders and non-binders for 3CLPro. This complementary screening approach is presented in detail in the section ‘Machine learning based compound screening’ below.

**Figure 1.**
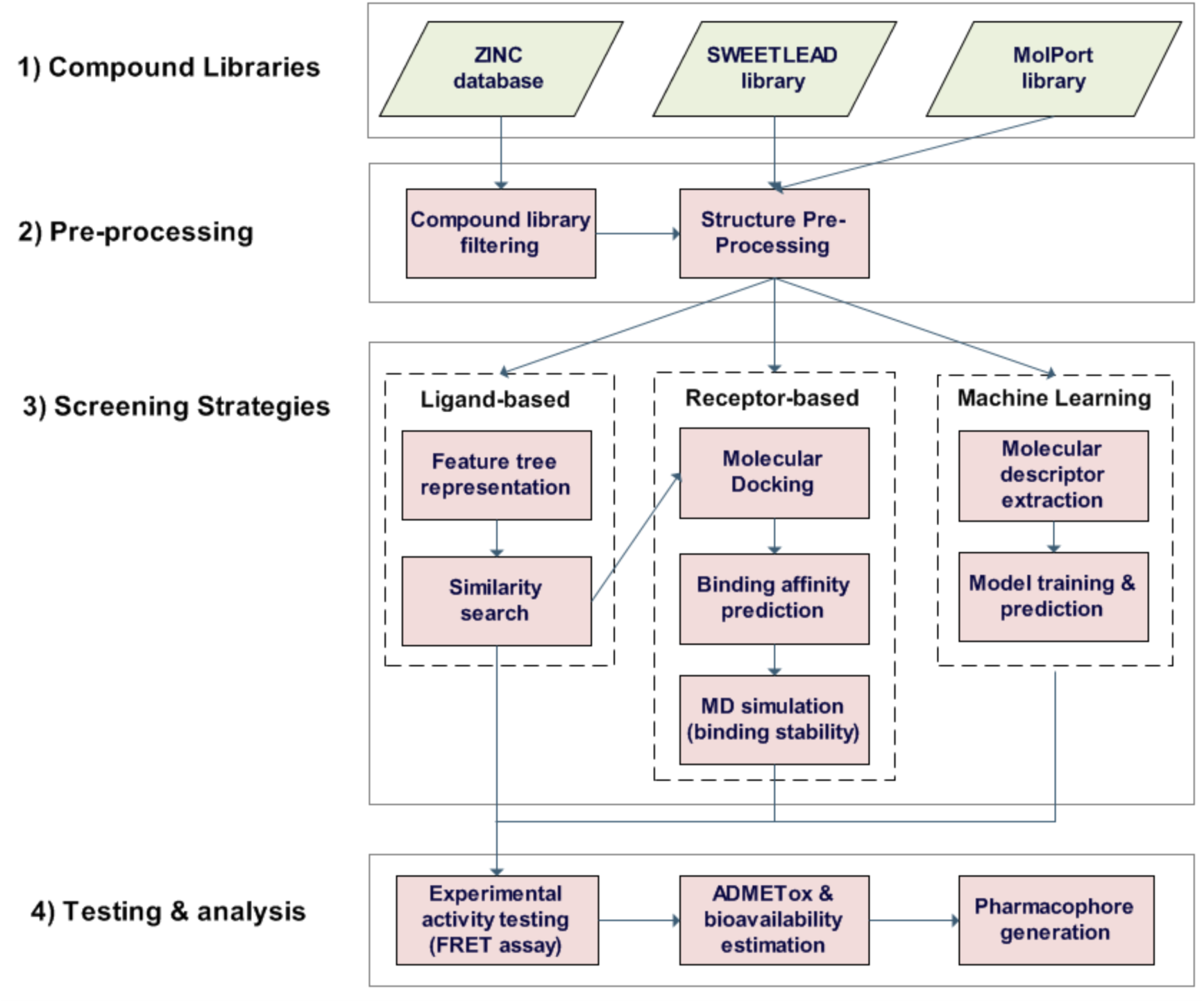
Overview of overall workflow for the screening procedure, starting with a compound library of > 1 billion compounds from the databases ZINC, SWEETLEAD and the MolPort. Due to the large size of the ZINC database, this compound repository was pre-filtered to focus on compounds that are commercially available and have drug-like chemical and ADMETox properties, and receptor-based screening was only applied after prior ligand-based screening (see section on ‘Ligand pre-processing and filtering’). More computationally expensive approaches relying only on receptor-based screening were only applied to the smaller SWEETLEAD and MolPort libraries, and machine learning based compound screening was only applied to the MolPort library (see sections on ‘Receptor-based screening using molecular docking’ and ‘Machine learning based compound screening’). A final selection of 95 top-ranked compounds derived from the *in silico* screening approaches was tested experimentally using a FRET assay to assess the 3CLpro inhibitory activity. Confirmed active compounds were characterized in terms of their known or computationally estimated ADMETox and bioavailability properties. Finally, pharmacophore models were generated for the active compounds.

The compounds prioritized using these *in silico* screening methods were then evaluated experimentally to determine the subset of stable binders and their 3CLpro inhibitory activity, using an *in vitro* Förster-type resonance energy transfer (FRET) assay. These experimental analyses are described in more detail in the section ‘*in vitro* assay for 3CLpro inhibitory activity assessment’.

### COMPUTATIONAL METHODS

#### Protein structure pre-processing

Three publicly available protein crystal structures for 3CLPro from the Protein Data Bank [28] were chosen for the molecular docking analyses and binding affinity estimation (PDB: 5R8T, 6YB7 and 6LU7). The structures 5R8T and 6YB7 were selected mainly due to their resolution (5R8T: 1.27 Å, 6YB7: 1.25 Å) and quality (R-free value: 0.208 for 5R8T and 0.192 for 6YB7), see additional quality assessments described below), whereas the structure 6LU7 was used additionally as a representation of the holo form of the protein in complex with an inhibitor (resolution: 2.16 Å, R-free value: 0.235), allowing us to compare docking results across different types of structures. Since many previously reported 3CLpro inhibitors are allosteric inhibitors [29–31], and allosteric inhibitors have been proposed to disrupt the 3CLpro dimer or slow the rate of processing significantly [29], we did not focus specifically on targeting the 3CLpro dimerization interface, but aimed to cover all binding pockets associated with already identified small molecule inhibitors. As an alternative approach, for the SARS-CoV version of 3CLpro, previous studies have also investigated peptide inhibitors designed specifically to target the dimerization [32, 33].

The receptor structures were pre-processed using the Schrödinger Maestro software (version 11.8.0.1.2, www.schrodinger.com) by adding hydrogens, generating protonation states, and optimizing hydrogen positions using the ‘Protein Preparation Wizard’ with default settings. The quality of the original and processed structures was assessed using the software tools Verify3D [34], WHATCHECK [35] and PROCHECK [36] as implemented in the software SAVES 5.0 (http://servicesn.mbi.ucla.edu/SAVES), confirming the suitability of the structures for docking simulations in terms of common quality control checks [37]. For structure files containing multiple chains, the chain with the highest Verify3D score was chosen for further analysis.

#### Ligand pre-processing and filtering

Small molecule compounds were obtained from the databases ZINC (version: ZINC15, May 2020, [22]), SWEETLEAD (version: 1.0, [23]), and the public library provided by the company MolPort (May 2020, www.molport.com). All ligands from the SWEETLEAD database were first preprocessed using the AutoDock ligand preparation script, and docked into the 3CLpro binding pocket using AutoDock-GPU [38]. For the ZINC database, in order to focus on compounds that are commercially available and have drug-like chemical and ADMETox properties, a filtering was first applied by downloading only compounds with the properties “drug-like”, “purchasable” (minimum purchasability = “Wait OK”) and reactivity = “clean”. This first-step filtering reduced the initial 1,276,766,435 substances to 898,838,573 retained substances. These filtered compounds were downloaded in SMILES-format, using the “ZINC-downloader-2D-smi.wget” script derived from the “Tranches” web-page on ZINC (https://zinc.docking.org/tranches/home).

The pre-selection of compounds from ZINC was then further filtered using a ligand-based similarity screening. For this purpose, a similarity assessment approach involving a graph-based molecular representation known as “feature trees” was applied, as implemented in the BioSolveIT Ftrees software (version 6.2). With this method the topological and physicochemical similarity of the library compounds to known small-molecule inhibitors for 3CLpro reported in the literature was scored (considering inhibitors for both the SARS-CoV and SARS-CoV-2 versions of the 3CLpro protein). Specifically, the literature-derived query compounds include the reported *SARS-CoV-2* 3CLpro inhibitors GC-376 [18], ebselen [39, 40] and baicalein [41], and the reported *SARS-CoV* 3CLpro inhibitors amentoflavone [42], hesperetine [43], pectolinarin [44] and dieckol [45]. Moreover, to further extend the search space of potential candidate inhibitor compounds, four additional query compounds for the feature tree search were included by adding the top-ranked compounds from the initial AutoDock-GPU screening (see above). All compounds from the ZINC library exceeding a minimum similarity threshold of 0.8 in the FTrees screen to these query compounds were retained for the subsequent molecular docking analyses.

For the screening of the MolPort compound library, due its association with our main provider for the commercial ordering of compounds and its smaller size (∼7.7 million compounds) compared to the ZINC library, we tested multiple alternative more extensive screening approaches without prior library filtering: (1) a screening approach relying purely on fast molecular docking approaches (see details in following section), (2) a screening approach relying purely on machine learning (see section on ‘Machine learning based compound screening’ below), and (3) a combination of molecular docking and ligand-similarity based screening using the software FTrees (following the same approach as for the ZINC database, but without prior database filtering, and focusing on the most potent available 3CLPro inhibitor, GC-376, as query compound; see following section for the description of the docking analyses).

Conformers for the collected compounds were generated using the OpenEye OMEGA software (version: 3.1.2.2, http://www.openeye.com) using the classic mode with default parameters.

#### Receptor-based screening using molecular docking

In order to obtain a robust ranking of compounds using molecular docking, the compound libraries prepared in the previous step (i.e. the filtered version of the ZINC database, and the unfiltered SWEETLEAD and MolPort databases) were docked into the 3CLpro binding pocket using three different approaches, AutoDock-GPU (an OpenCL and Cuda accelerated version of AutoDock4.2.6, [38]), OpenEye HYBRID (version 3.4.0.2, [46–48]), and LeadIT/FlexX (version 2019.Nov.2-Ubuntu-18.04-x64, [49]). To use the available computing time efficiently and only consider compounds with high relative ranking scores for all docking approaches for further analysis, the two most time-efficient docking methods AutoDock-GPU (with the parameter ‘nrun’ for the thoroughness of the search space exploration set to 100, and default parameters otherwise) and OpenEye HYBRID (with default docking parameters) were first run on the pre-selected compound libraries, and only compounds with higher than average scores from the AutoDock-GPU and OpenEye HYBRID screens were also docked using FlexX (with default docking parameters), including a subsequent estimation of the binding affinity for the top 30 docking poses using the LeadIT/HYDE approach [50]. The docking methods are semi-flexible, taking ligand flexibility into account; however, as a limitation in contrast to recent supercomputer-based screening approaches, which address receptor flexibility using high-resolution ensembles of many 3CLpro receptor conformations [25], ensemble docking and enhanced sampling MD [24], the assessment of receptor flexibility in the present study was restricted to the MD simulations applied to the top-ranked compounds derived from the docking analyses (see section on ‘Molecular dynamics simulations’).

For the MolPort library, we compared both a screening approach relying purely on molecular docking of the unfiltered library, and a joint docking and ligand similarity-based scoring approach, which filtered the compounds by including only those reaching a minimum similarity threshold of 0.8 in the ligand-based screening using FTrees (see previous section). Finally, the list of compounds docked with each method was ranked and sorted according to the sum of ranks across the scores for all methods. Only top-ranked compounds achieving consistently high docking scores for the three pre-processed 3CLpro protein structures (PDB: 5R8T, 6YB7 and 6LU7) were used for experimental validation. Visualizations of the top-scoring docked compounds were generated in the software Chimera (version 1.12, [51]) and binding interactions were analyzed using PoseView (version 1.1.2, [52]). A list of all used software tools and their versions, as well as scripts including the commands and parameters for the virtual screening analyses are provided on GitHub (https://github.com/eglaab/3clpro).

#### Molecular dynamics simulations

To assess the ligand binding stability for the top-ranked compounds derived from the docking simulations in the 3CLpro binding pocket, molecular dynamics (MD) simulations were carried out using a GPU-accelerated version of the software NAMD (Git-2020-07-30 for Linux-x86_64-multicore-CUDA, [53]). Topology and force field parameters for the MD simulations were assigned from the CHARMM36 protein lipid parameter set [54, 55] for the receptor and from the CHARMM General Force Field (CGenFF) parameter set [56] for the ligands. The step size was 2 fs, and the simulation was performed for a duration of 20 ns. Video animations of the simulations were created by applying the “MD Movie” function in the software Chimera (version 1.12, build 41623, [51]) to the trajectory data for the protein-ligand complexes derived from NAMD. The detailed configuration settings used for NAMD and exemplary video animations of MD simulations for confirmed inhibitors are provided on GitHub (https://github.com/eglaab/3clpro).

#### Machine learning based compound screening

As an alternative to conventional receptor- and ligand-based screening strategies, we also tested a compound ranking approach using machine learning. For this purpose, quantitative properties for the investigated compounds were determined by calculating molecular descriptors as implemented in the R software package “rcdk” (version 3.5.0) [57] using SMILES compound representations [58] as input. First, molecular descriptors were computed for a previously published training set collection of small molecules containing both previously reported SARS-CoV or SARS-CoV-2 3CLPro inhibitors (486 compounds, used as positive set) and compounds reported not to bind to 3CLPro (700 compounds, used as negative set) [59]. The covered classes of descriptors in the package “rcdk” include topological descriptors (e.g. TPSA, Wiener index, Kier-Hall Smarts descriptor), geometrical (e.g. gravitational index, Petitjean shape index, moment of inertia), electronic (e.g. CPSA, H-bond donor count, H-bond acceptor count), constitutional (e.g. rotatable bond count, acidic group count, weight) and hybrid descriptors (WHIM and BCUT descriptors; see the documentation of the rcdk software package to retrieve the complete list of more than 200 descriptors).

A classification model using the Random Forest algorithm as implemented in the R software package “randomForest” (version 4.6-14, [60]) was then built on this training data of molecular descriptors for the positive and negative compound sets, using 250 decision trees and default settings otherwise. This model was applied to all compounds from the MolPort library, by computing the descriptors for each compound as input to the model and generating probabilistic predictions. Finally, the predicted probability for each compound to belong to the positive set of 3CLPro inhibitors was used to rank the compounds, and the top 20 compounds with the highest probabilities were selected for *in vitro* testing. A script for the machine learning analyses has been made available on GitHub (https://github.com/eglaab/3clpro).

### EXPERIMENTAL METHODS

#### In vitro assay for assessing the 3CLpro inhibitory activity of the compounds

##### Assay principle

Förster resonance energy transfer (FRET) assays can be applied in various configurations in cells [61] and in reporter systems [62] to detect molecular proximity and with the proper biosensor also activity. Here we employed an *in vitro* FRET assay for the evaluation of the candidate small molecule inhibitors for 3CLpro obtained from the *in silico* screening approaches. We used a substrate peptide of 3CLpro labelled with a fluorescent dye (Dabcyl) and an acceptor-quencher (Edans) at the N- and C-terminus, respectively. The substrate peptide does not fluoresce in the uncleaved state, when the quencher blocks the fluorescence of the dye. When 3CLpro cleaves the substrate, the fluorescence of the dye is de-quenched and an emission signal is observed. The inhibitor blocking the activity of 3CLpro prevents FRET-peptide cleavage. Thus, in the presence of an inhibitor a lower fluorescence signal is observed [63]. A similar FRET-based assay has recently been applied to test candidate 3CLpro inhibitor compounds derived from the lead optimization of a prior week screening hit, and some of these compounds were also successfully confirmed to inhibit infectious SARS-CoV-2 replication in Vero E6 cells [21, 27]. Moreover, after the completion of our experimental studies, an optimized assay for high-throughput screening of 3CLpro using a new fluorogenic substrate became available [20]. This assay has been used to identify two irreversible 3CLpro inhibitors, azanitrile (K_i_ = 24.0 nM) and pyridyl ester (K_i_ = 10.0 nM) [20], as candidates for further development and provides a more scalable methodology than the one applied in the present study. Thus, for follow-up *in vitro* screening studies of 3CLpro inhibitors, more recently developed assays may be preferred over the assay with lower throughput employed here.

##### Method

The assays were performed following the guidelines of the 3CL Protease assay Kit (#79955, BPS Biosciences, San Diego CA, USA). The inhibitory effect on 3CLpro was assessed using the substrate peptide Dabcyl-KTSAVLQSGFRKM-E(Edans)-NH2, obtained from Biosyntan. The assays were run on a 384-well plate (black, low volume, round bottom; Corning #4514) and the reaction volume per well was 20 µl. Briefly, all candidate inhibitors were dissolved in DMSO and three-fold or five-fold diluted in the assay buffer. The known 3CLpro inhibitor and reference compound GC-376 was dissolved in water and diluted from 100 µM. To the dilution series of inhibitors, 3CLpro (100 ng/reaction) was added and incubated for 30 min at room temperature (RT). Then 50 µM of substrate peptide was added. The reaction mix was incubated at RT overnight with the plates sealed. On the next day, fluorescence intensity was measured at excitation wavelength 360 ± 10 nm and emission 460 ± 20 nm using Clariostar (BMG Labtech, Germany) plate reader. 3CLpro used in the assay was N-terminally MBP-tagged (MW – 77.5 kDa) and obtained from BPS Biosciences (#100707). The data were fit into a log c(inhibitor) vs response three-parameter dose-response model using the software Prism (GraphPad). IC_50_ value estimates were determined individually for each replicate, and the mean and standard error of the mean (SEM) were calculated. Importantly, many of the assessed compounds were polyaromatic natural products, which exhibited interfering fluorescence or absorbance in our FRET-based assay. Therefore, out of 96 compounds tested in the 3CLpro activity assay, 12 compounds were filtered out from further analysis (highlighted in blue color in the lists of tested compounds in the Supporting Information). Assays involving principles of time-resolved FRET may be suitable for future follow-up screening of compounds with background fluorescence [64, 65].

## RESULTS AND DISCUSSION

### In silico screening results and final compound selection for in vitro testing

For the experimental testing of candidate 3CLPro small molecule inhibitors, 86 commercially available compounds selected from the top-scoring candidates derived from the different receptor-, ligand- and machine learning based screening approaches (see Methods), which had passed all prior *in silico* filtering stages, were ordered from the company MolPort. The complete list of tested screening compounds, including common compound identifiers and characteristics (SMILES, molecular weight, logP) is provided in the Supporting Information. The 86 top-ranked compounds are subdivided into 18 compounds derived from the virtual screening of the ZINC and SWEETLEAD libraries (Table S2 in the Supporting Information; see Methods for the prefiltering and screening approaches), 30 compounds derived from the screening of the MolPort library using combined ligand similarity screening and molecular docking (Table S3), 18 compounds from the screening of the MolPort library using combinations of docking algorithms only (Table S4), and 20 compounds from the purely machine learning derived compound ranking for the MolPort library (Table S5). In addition to these screening-derived compounds, the known inhibitor GC-376 [18] was included as a reference compound, and the following commercially available compounds previously reported as SARS-CoV-2 or SARS-CoV 3CLPro inhibitors in the literature, were also acquired for experimental inhibitory activity assessments: Ebselen [39, 40], baicalein [41], amentoflavone [42], hesperetine [43], pectolinarin [44], luteolin [42], quercetin [42], pristimerin [66], and 1-hydroxypyridine-2-thione zinc [67]. For the compounds confirmed as binders (see Table 1), PAINS and Brenk’s substructure alerts [68, 69] were checked to determine frequent hitters (promiscuous compounds, see Table 2). GC-376 is a prodrug that converts into the peptide aldehyde GC-373 [70]; hence, the active compound is GC-373, and the inhibitor is therefore referred to as “GC-376 / GC-373” in the following. Example video animations of the MD simulations which were applied to the top-ranked docking-derived compounds to confirm the stability of the binding are provided online together with the configuration settings used for the MD simulation software NAMD (https://github.com/eglaab/3clpro).

**Table 1:**
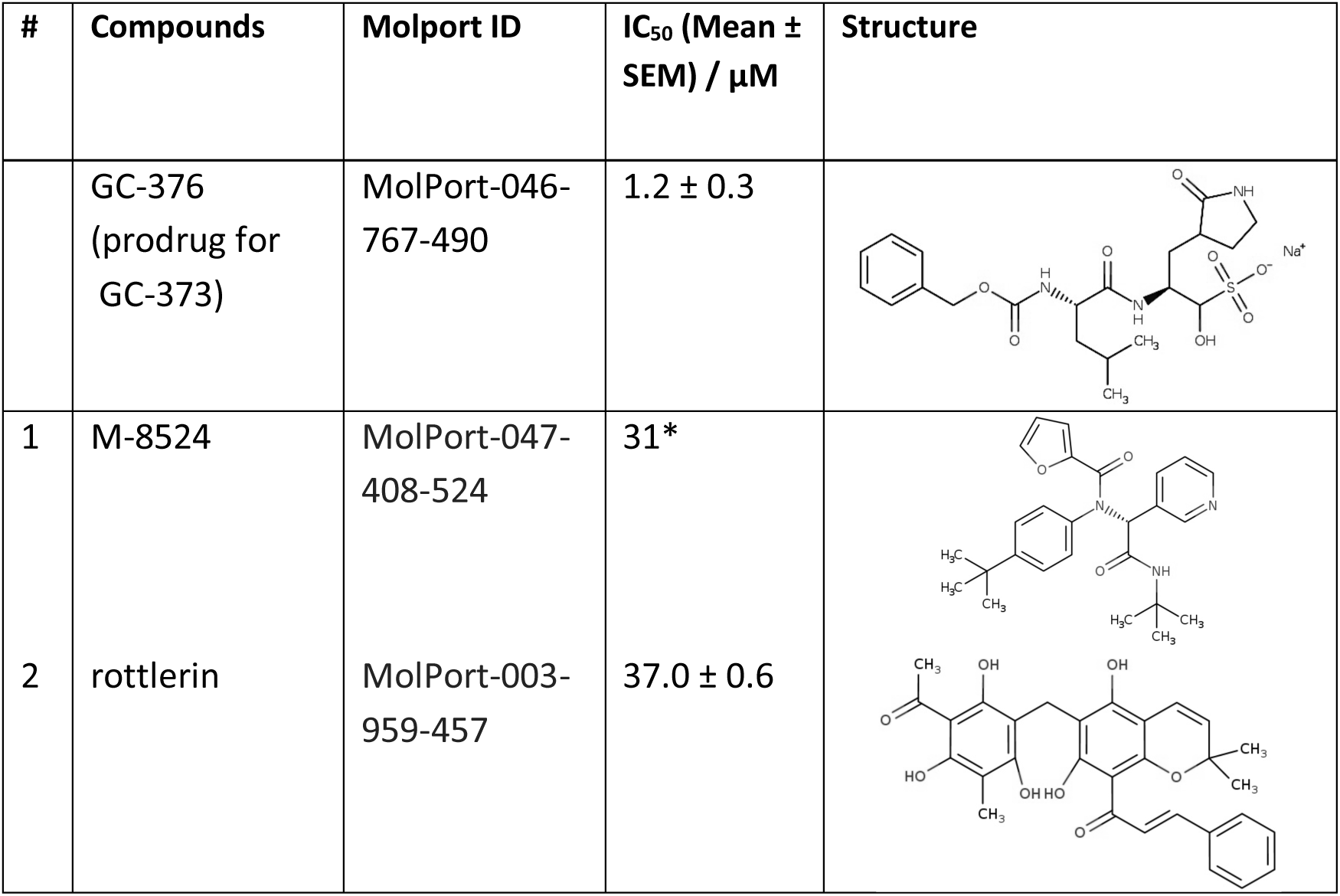
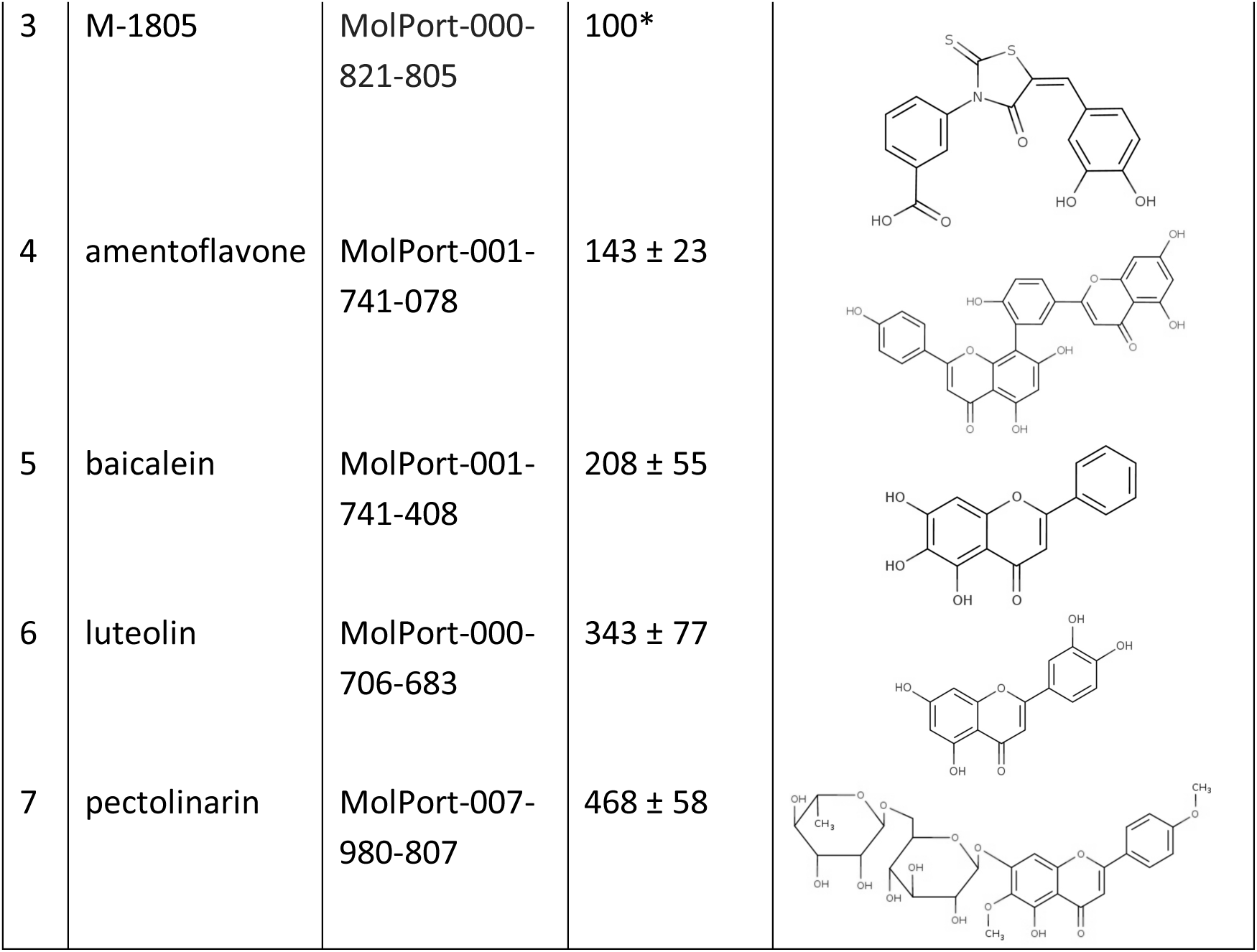
Hits from 3CLpro screening assay with IC^50^ values. *For the compounds no. 1 and 3 only a limited amount of the reagent could be acquired, and therefore no replicate runs could be performed for these specific compounds.

| # | Compounds | Molport ID | $IC_{50}$ (Mean ± SEM) / $\mu M$ | Structure |
| --- | --- | --- | --- | --- |
| | GC-376<br>(prodrug for GC-373) | MolPort-046-767-490 | $1.2 \pm 0.3$ | |
| 1 | M-8524 | MolPort-047-408-524 | 31* |  |
| 2 | rottlerin | MolPort-003-959-457 | $37.0 \pm 0.6$ | |
| 3 | M-1805 | MolPort-000-821-805 | 100* |  |
| 4 | amentoflavone | MolPort-001-741-078 | 143 ± 23 |  |
| 5 | baicalein | MolPort-001-741-408 | 208 ± 55 |  |
| 6 | luteolin | MolPort-000-706-683 | 343 ± 77 |  |
| 7 | pectolinarin | MolPort-007-980-807 | 468 ± 58 |  |

**Table 2:** Estimated drug- and lead-likeness and bioavailabilty scores (bioavailability is low = below 0.25, medium = between 0.25 to 0.75, high = above 0.75) of the confirmed 3CLpro inhibitors as estimated using the SwissADME software (more detailed information on the specific score violations and alerts as well as other physicochemical and ADME estimations can be found in the Supporting Information). Passed filters are highlighted in green color, failed tests are marked in red color.

| Compounds | GC-376 | M-8524 | rottlerin | M-1805 | amento-flavone | baicalein | luteolin | pectolinarin |
| --- | --- | --- | --- | --- | --- | --- | --- | --- |
| Druglikeness (Lipinski) | No (2 violations) | Yes | Yes (1 violation) | Yes | No (2 violations) | Yes | Yes | No (3 violations) |
| Druglikeness (Ghose) | No (1 violation) | Yes | No (2 violations) | Yes | No (2 violations) | Yes | Yes | No (4 violations) |
| Druglikeness (Veber) | No (2 violations) | Yes | No (1 violation) | No (1 violation) | No (1 violation) | Yes | Yes | No (1 violation) |
| Druglikeness (Egan) | No (1 violation) | Yes | No (1 violation) | No (1 violation) | No (1 violation) | Yes | Yes | No (1 violation) |
| Druglikeness (Muegge) | No (1 violation) | Yes | No (1 violation) | No (1 violation) | No (3 violations) | Yes | Yes | No (4 violations) |
| Bioavailability Score | low | medium | medium | low | low | medium | medium | low |
| PAINS | 0 alerts | 0 alerts | 0 alerts | 2 alerts | 0 alerts | 1 alert | 1 alert | 0 alerts |
| Brenk | 1 alert | 0 alerts | 1 alert | 3 alerts | 0 alerts | 1 alert | 1 alert | 0 alerts |
| Lead-likeness | No (2 violations) | No (3 violations) | No (2 violations) | No (1 violation) | No (2 violations) | Yes | Yes | No (2 violations) |

### *In vitro* testing results for candidate 3CLpro inhibitors

In total, 95 selected compounds were assessed to determine their *in vitro* 3CLpro inhibitory activity, including the 86 candidate compounds identified from the *in silico* analyses and 9 candidate compounds derived from the literature, as described above. GC-376 was used as a positive control for 3CLpro inhibitory activity. The IC_50_ value of GC-376 was in the low micromolar range (0.18 to 0.98 µM), matching approx. with previously reported IC_50_ values for GC-376 (0.19 μM) and its structurally similar metabolized form, GC-373 (0.4 μM) [70] (see Supporting Information for the dose-response curves). The Z’ value [71], used as a measure of assay robustness, ranged from 0.5 to 0.8, indicating the suitability of the assay for compound screening in a high-throughput format. Importantly, the main purpose of the activity assay is to obtain a robust relative ranking of ligands in terms of their inhibitory activity rather than high precision absolute activity estimates.

Overall, 7 out of the 95 tested compounds were confirmed as active, with a lowest IC_50_ of 31 µM for the screening compound M-8524 (CID 46897844, see Table 1). Considering the different conducted screening analyses separately, each type of strategy uniquely identified at least one of the confirmed inhibitors: (1) The combination of molecular docking and ligand similarity screening on the ZINC database identified the inhibitor rottlerin among 18 compounds tested, (2) the screening of the MolPort database using fast docking approaches detected another inhibitor (compound M-1805) out of 18 tested candidates, and (3) the screening of the MolPort library using machine learning identified a further inhibitor out of 20 assessed compounds. The complete list of active compounds, including their names, vendor ID, IC_50_ values and 2D structure representations are shown in Table 1. Detailed assay validation results and dose-response curves for these inhibitors are provided in the Supporting Information.

### ADMETox and bioavailability properties of the 3CLpro screening assay hits

While the identified inhibitors did not display a higher activity than the currently available most potent inhibitor GC-376 / GC-373, used as reference compound, multiple of the confirmed actives are natural compounds with favorable bioavailability and ADMETox (Absorption, Distribution, Metabolism, Excretion and Toxicity) properties, that may provide a basis for follow-up structure design optimization efforts or further ligand-based similarity screening in independent compound databases.

To compare all actives in terms of ADME and physicochemical characteristics, corresponding parameters of the confirmed inhibitors were estimated computationally using the software SwissADME (see Table 2 for a summary of key features and Figures S4 to S11 in the Supporting Information for a more comprehensive overview of parameters). This analysis suggested that a subset of compounds, including M-8524, baicalein and luteolin, fulfil multiple typical properties of drug-like, orally bioavailable compounds in terms of the drug-likeness scores proposed by Lipinksi et al. [72], Ghose et al. [73], Veber et al. [74], Egan et al. [75], Muegge et al. [76], and have Abbott Bioavailability Scores above 0.5 [77]. Moreover, some of these compounds also pass all or most PAINS and Brenk’s filters [68, 69] or display lead-like properties (250 g/mol ≤ molecular weight ≤ 400 g/mol, xlogP ≤ 3.5, num. rotatable bonds ≤ 7) [78], potentially providing a basis for further structural and chemical optimization. By contrast, the reference compound GC-376 is not drug-like or lead-like according to these scores and has a low bioavailability score (0.17, see Table 2 and Supporting Information).

Apart from the computationally estimated ADME properties, prior information on bioavailability and ADMETox characteristics was available for some of the natural products among the confirmed inhibitors. The natural compound with the lowest IC_50_ value, rottlerin, is derived from the powder that covers the capsules of the tree *Mallotus phillippinensis* (also known as Indian Kamala, Rottlera tree, or monkey-faced tree) [79], which mainly occurs in Asia, Afghanistan and Australia. The powder has traditionally been used as antiparasitic agent against the tapeworm in India for multiple centuries, providing a first indication of favorable safety properties. In mouse model studies, rottlerin was investigated as a preventive or adjuvant supplement for pancreatic cancer [80], as a protective agent against cisplatin-induced nephrotoxicity [81], and as a potential medication in a psoriasis model, where it was well tolerated with no significant changes in bodyweight [82]. Tissue bioavailability and pharmacokinetic analyses *in vitro* and *in vivo* suggest that rottlerin is efficiently absorbed in cells and tissues [80].

The natural product with the second lowest IC50 is amentoflavone, which had previously already been reported to inhibit the 3CLpro protein from SARS-CoV (the virus from the 2002-2004 SARS outbreak) [42]. However, a later study suggested that the binding reaction results from a non-specific aggregate formation or complexation, because no effective inhibition was observed in the presence of 0.01% Triton X-100 [44]. Amentoflavone is a biflavonoid contained in several plant species including *Ginkgo biloba*, *Hypericum perforatum* (St. John’s wort), *Xerophyta plicata* and *Chamaecyparis obtusa*. No detailed prior toxicological studies of amentoflavone could be identified, but several bioactivities have been reported in the biomedical literature, including anti-viral effects [83, 84].

Among the other natural compounds confirmed to bind to SARS-CoV-2 3CLpro, baicalein displays some of the most favorable drug-likeness scores (see Table 2). This flavonoid is found in the dried roots of the plant *Scutellaria baicalensis (S. baicalensis) Georgi* [85], and was previously already reported to display an antiviral activity against SARS-CoV *in vitro* [86] and to inhibit the SARS-CoV-2 3CL protease *in vitro* [41]. The use of baicalein containing roots was widespread in traditional Chinese herbal medicine over centuries, and a wide range of pharmacological activities have been identified. These include antiviral activities were reported for further virus species apart from SARS corona viruses, including dengue virus [87, 88], H5N1 influenza A virus [89], and Japanese encephalitis virus [90].

Luteolin, a structurally similar compound to baicalein, also displayed similar estimated ADME properties in terms of favorable drug-likeness and bioavailability scores (see Table 2), and an inhibitory effect of luteolin on SARS-CoV 3CLpro was observed in a previous study using a FRET assay (IC50 = 20.2 µM) [42]. Regarding the bioavailability of luteolin, oral administration of pure luteolin in rats (14.3 mg/kg) has been shown to result in a limited peak plasma concentration of 1.97 ± 0.15 μg/mL, which was increased to 8.34 ± 0.98 μg/mL when delivering the compound in peanut hull extracts [91]. Luteolin is contained in small quantities in both medicinal plants and commonly consumed vegetables and spices (e.g. broccoli, thyme, pepper, and celery), and displays no mutagenic effects in the Ames test in contrast to other flavonoids [92], such as quercetin. Interestingly, quercetin, which only differs structurally from luteolin by the position and number of hydroxyl groups, was also reported to bind to SARS-CoV 3CLpro [42], but with a slightly higher IC50 than luteolin (IC_50_ = 23.8 µM). When testing quercetin as a candidate inhibitor of SARS-CoV-2 3CLpro in the FRET analyses reported here, it did however not show any activity. This suggests that the hydroxyl group positions in flavonoids are an important determinant of their activity for SARS-CoV-2 3CLpro.

Finally, the last natural product confirmed to bind to SARS-CoV-2 3CLpro is pectolinarin. It was previously already found to block the enzymatic activity of SARS-CoV 3CL protease according to a FRET protease assay, using a similar approach as in the study presented here [44]. The compound is isolated from plants of the genus *Cirsium*. Specifically, the plant *Cirsium setidens*, which contains pectolinarin as a major active compound, has been used in traditional Korean medicine against hypertension, hemostasis, hematoma and hematuria [93]. Regarding prior pharmacokinetics studies for pectolinarin, after oral administration of 6 mL/kg *Cirsium japonicum DC* extract in rats, the maximum plasma concentration of the component pectolinarin was 877 ±97 ng/mL, reached after 5 min. [94]. However, pectolinarin displays low drug- and lead-likeness and bioavailability scores in the computational estimations (see Table 2).

### Pharmacophore model for 3CLpro small molecule inhibitors

In order to generate a pharmacophore model that describes characteristic features of 3CLpro small molecule inhibitors, we investigated the interactions of the identified inhibitors with residues in the receptor binding pocket using the best scoring docking pose for each compound according to the binding affinity estimation by the software HYDE. For the reference inhibitor GC-376 / GC-373 used for comparison, the relevant pose was derived from a crystal structure which contains this compound in complex with 3CLPro (PDB ID: 7D1M). Similarly, for compound baicalein a crystal structure of the complex with 3CLpro was already available (PDB ID: 6M2N) and used to extract the binding pose. Figure 2 shows the 2D representations of the estimated hydrogen bond interactions (dotted lines) and hydrophobic contacts (green lines) for each inhibitor in the 3CLPro binding pocket as determined using the software PoseView [52].

**Figure 2.**
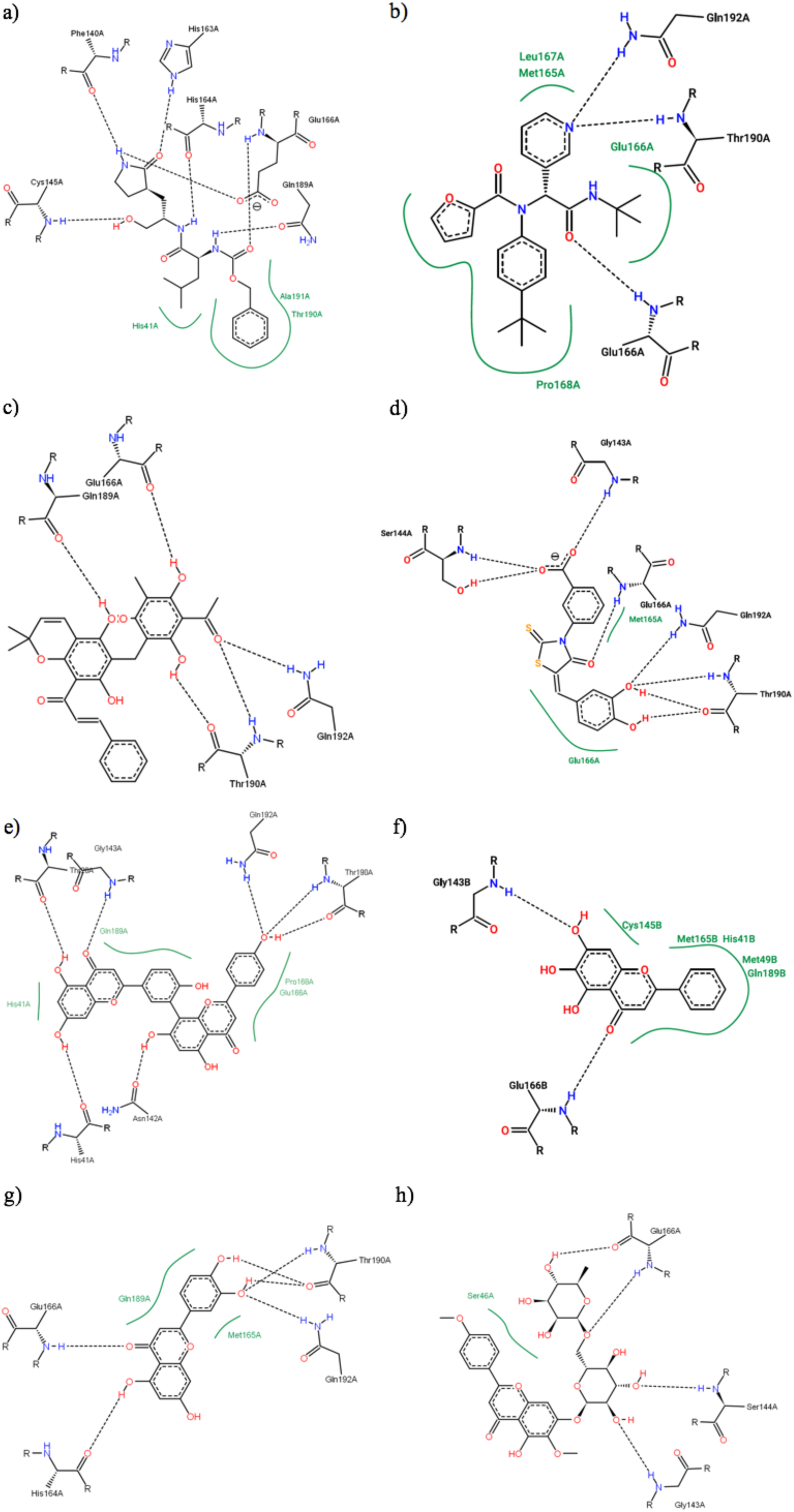
2-dimensional representations of the hydrogen bond interactions (dotted lines) and hydrophobic contacts (green lines) between confirmed small molecule inhibitors and residues in the 3CLPro binding pocket: a) GC-373, b) M-8524, c) rottlerin, d) M-1805, e) amentoflavone, f) baicalein, g) luteolin, h) pectolinarin. Plots were generated using the software PoseView. The inhibitor GC-376 / GC-373 is shown in the pose derived from a crystal structure of 3CLPro in complex with this compound (PDB ID: 7D1M). All other sub-figures represent the best scoring docking poses for the corresponding compounds according to the HYDE binding affinity estimation score.

An overview of the binding pocket residues involved in these protein-ligand interactions is provided in Table 3, highlighting all interactions shared by at least two compounds in color, and representing identical residues by identical colors. Most hydrogen bond interactions and hydrophobic contacts with binding pocket residues are shared by multiple ligands, suggesting that they reflect a common binding mode and a characteristic signature of interactions that favor stable ligand binding. The residues most frequently participating in these protein-ligand interactions are Glu166 (involved in hydrogen bonds for 8 inhibitors, and in hydrophobic contacts for 3 inhibitors), Thr190 (H-bonds: 5, Contacts: 1), Gln189 (H-bonds: 2, Contacts: 3) and Gln192 (H-bonds: 5). Interestingly, the two residues His41 and Cys145 which have previously been confirmed to be critical for catalysis [95] are also involved in hydrogen bond interactions with some of the inhibitors (His41 for amentoflavone; Cys145 for GC-376 / GC-373) and hydrophobic contacts (His41 for GC-376 / GC-373, amentoflavone, and baicalein; Cys145 for baicalein), and these compounds may directly interfere with the catalytic mechanism. Overall, both the shared interactions (see Table 3) and the graphic overlay of the docked or crystal structure derived ligand conformations (see Figure 3) suggest that the confirmed inhibitors use a similar binding mode to engage the 3CLpro binding pocket.

**Figure 3.**
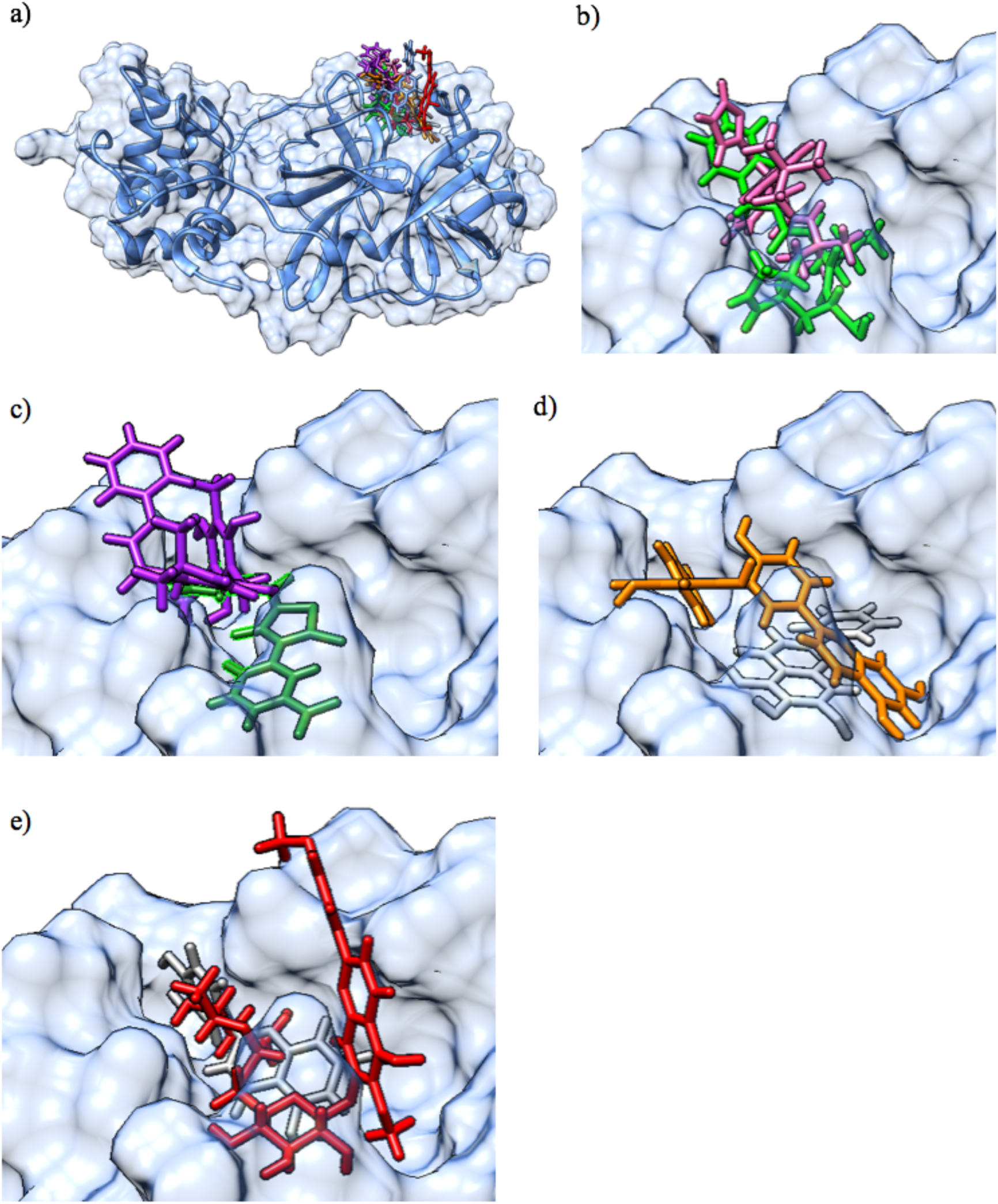
a) Molecular surface representation of the 3CLpro protein with an overlay of all confirmed actives in the binding pocket (top right) in their best scoring docking conformations (only for GC-376 / GC-373 and baicalein, crystal structures of the protein-ligand complex were available and used instead of molecular docking derived conformations). The 3CLpro secondary structure is displayed as a blue ribbon, surrounded by the solvent excluded molecular surface representation in transparent blue, and with the colored ligands shown in a stick representation. b) Larger scale representation of the overlaid pair of ligands with the lowest IC_50_ values: GC-376 (light green) and M-8524 (pink) in the 3CLpro binding pocket (top right part of figure 3 a); c) overlaid pair of ligands rottlerin (purple) and M-1805 (dark green); d) overlaid pair of ligands amentoflavone (orange) and baicalein (light gray); e) overlaid pair of ligands luteolin (dark gray) and pectolinarin (red).

**Table 3.** Overview of hydrogen bond interactions (2^nd^ column) and hydrophobic contacts (3^rd^ column) between the inhibitors and residues in the 3CLPro binding pocket (inferred by applying the PoseView software with default settings). The residue 3-letter codes and numbers correspond to the 3CLPro amino acids involved in the interaction type indicated in the column header (residues are sorted by increasing position in the 3CLpro amino acid sequence). All residues shown in color are involved in binding interactions for at least two different inhibitors, where identical colors represent identical residues. These shared interactions across multiple ligands may represent critical features that favor the binding of small molecules in the 3CLPro binding pocket.

| Compounds | H-bonds | Contacts |
| --- | --- | --- |
| GC-376 / GC-373 | Phe140, Cys145, His163, His164, Glu166, Gln189 | His41, Thr190, Ala191 |
| rottlerin | Glu166, Gln189, Thr190, Gln192 | - |
| M-8524 | Glu166, Thr190, Gln192 | Met165, Glu166, Leu167, Pro168 |
| M-1805 | Ser144, Gly143, Glu166, Thr190, Gln192 | Met165, Glu166 |
| amentoflavone | His41, Asn142, Gly143, Thr190, Gln192 | His41, Glu166, Pro168, Gln189 |
| baicalein | Glu166, Gly143 | His41, Met49, Cys145, Met165, Gln189 |
| luteolin | His164, Glu166, Thr190, Gln192 | Met165, Gln189 |
| pectolinarin | Gly143, Ser144, Glu166 | Ser45 |

Next, we created dedicated pharmacophore models for each of the inhibitors using the structure-based pharmacophore design approach in the software LigandScout [96]. Two example pharmacophore models are shown in Figure 4 for the reference compound GC-376 / GC-373 (Figure 4 a), and the newly identified inhibitor rottlerin (Figure 4 b), respectively. These two compounds occupy a similar space in the binding pocket, like most identified inhibitors in their docked binding conformation (see Figure 3), but engage at least partly in different hydrogen bond and hydrophobic interactions with the residues in the pocket. Thus, by merging the pharmacophore models for these and all other confirmed inhibitors into a single model, further virtual screening and structural design optimization efforts may exploit the unified model to identify new inhibitors with suitable steric properties that engage in more energetically favorable combinations of interactions than those covered by the already identified actives. Therefore, the pharmacophore models for the individual ligands were integrated by aligning the structures and their associated pharmacophores by receptor-derived reference points and interpolating the overlapping features, using the default settings in the LigandScout software. In contrast to the models for the individual inhibitors, the resulting merged pharmacophore model does not lend itself to direct interpretation, due to its larger coverage of potential binding interactions that can be exploited by a ligand. However, its more comprehensive coverage of potential interactions provides a computational resource for further screening studies on independent compound databases that is more informative than individual structures of protein-ligand complexes. The merged pharmacophore model has therefore been made available in the “Compressed Pharmaceutical Markup Language” (PMZ) format on GitHub (https://github.com/eglaab/3clpro). The reader should note that this ligand-based pharmacophore model is specific to the inhibitors identified in the present project, and should therefore ideally be considered in combination with other published pharmacophore models exploiting additional information sources, e.g. a recently proposed receptor-based pharmacophore definition for 3CLpro that combines information from molecular simulations and crystallographic studies [25].

**Figure 4.**
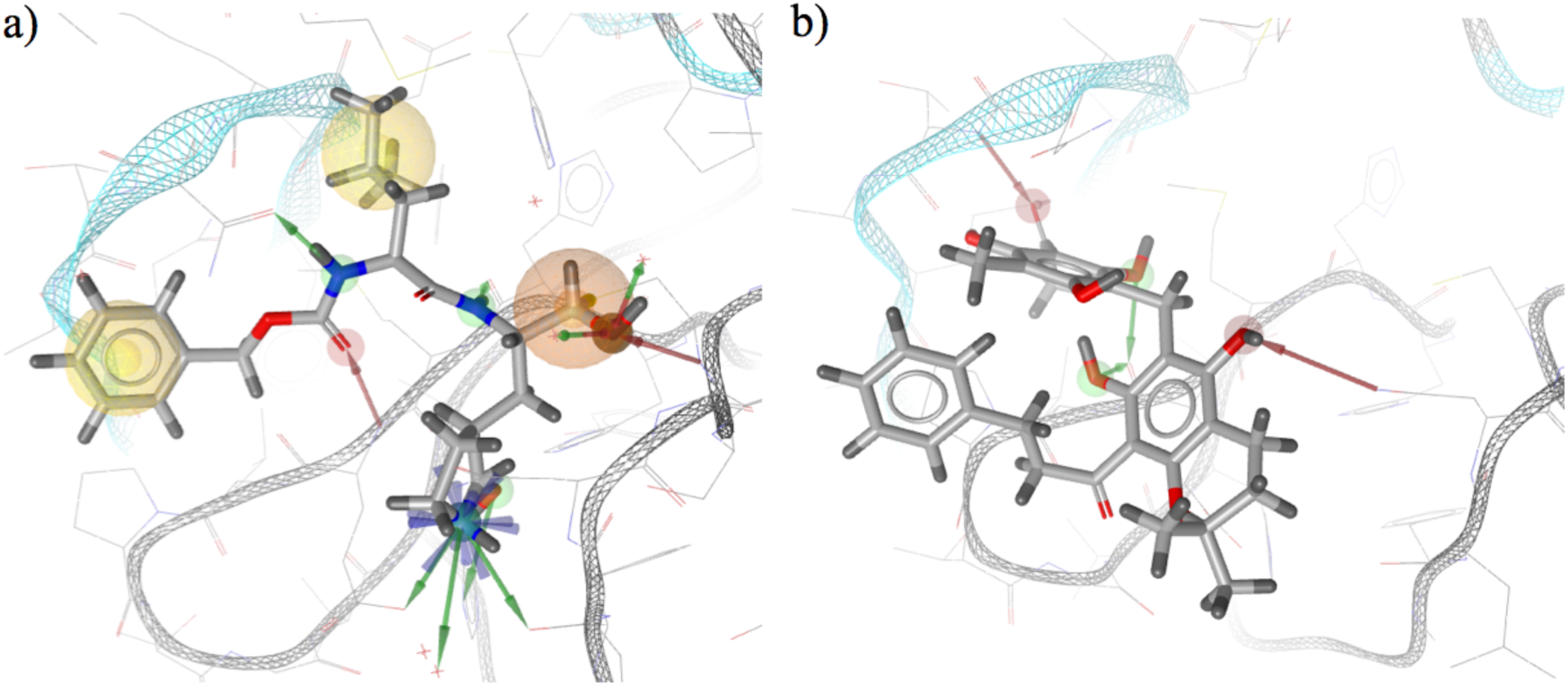
a) Pharmacophore model for 3CLpro derived from the reference inhibitor GC-376 / GC-373 using the software LigandScout (conformation obtained from the PDB crystal structure 7D1M after pre-processing with the software Schrödinger Maestro, see Methods). The yellow spheres represent hydrophobic interactions, the red spheres and arrows correspond to hydrogen bond acceptors, and the green spheres and arrows to hydrogen bond donors. b) Pharmacophore model for 3CLpro derived from the inhibitor rottlerin using the software LigandScout (conformation obtained from the best scoring pose in molecular docking according to the HYDE scoring approach, see Methods). While the comparison of rottlerin and GC-376 / GC-373 suggests that they occupy a similar space in the 3CLpro binding pocket and share a hydrogen bond interaction with the residue Glu166, most interactions are distinct. This may open up possibilities for the screening of new compounds that exploit interactions from the pharmacophore models of both the GC-376 / GC-373 and rottlerin ligand.

## CONCLUSIONS

The SARS-CoV-2 3CLpro protein is one of the main drug targets of interest for COVID-19, due to its critical role in viral replication, the availability of multiple high quality protein crystal structures and the prior identification of first small molecule inhibitors as a basis to computationally screen for inhibitors with improved inhibitory activity, bioavailability and ADMETox characteristics.

The combined computational and experimental analyses presented here reveal new natural and synthetic compounds inhibiting 3CLpro with micromolar activity, e.g. rottlerin and M-8524, and provide a pharmacophore model that describes key structural and chemical properties of active compounds. Both this pharmacophore and the confirmed hits provide a resource to support follow-up efforts on the structural design of more potent and selective 3CLpro inhibitors and similarity-based inhibitor screening in additional compound databases. The pharmacophore analyses suggest that many of the 3CLpro inhibitors share hydrogen bond and hydrophobic interactions with binding pocket residues, including interactions with the two residues His41 and Cys145, that have previously been confirmed to be critical for catalysis.

Apart from the new inhibitors identified through the screening of the ZINC, SWEETLEAD and MolPort databases, we also assessed compounds previously proposed as SARS-CoV 3CLPro inhibitors in the literature in terms of their SARS-CoV-2 3CLPro inhibitory activity, confirming amentoflavone, baicalein, luteolin and pectolinarin as actives (albeit with lower affinities then GC-376 / GC-373, rottlerin and M-8524, among others), and rejecting several other candidate compounds as actives (in the lists of tested compounds in the Supporting Information, the confirmed compounds are all marked with a star symbol; the compounds which could not be assessed in the assay due to high background fluorescence are marked in blue, and all other compounds were invalidated). Analyses of ADMETox and physicochemical parameters of the confirmed hit compounds show that these compounds differ significantly with regard to relevant known or computationally estimated properties, and that a subset of the hits displays favorable bioavailability and safety characteristics in terms of the prior knowledge available.

Regarding the computational methodological aspects of the project, as part of the virtual screening we have tested different screening approaches, including the integration of molecular docking and ligand similarity screening, the combined application of fast docking approaches without prior compound library filtering, and the use of machine learning for screening. The observation that each of these approaches uniquely identified at least one of the experimentally confirmed inhibitors suggests that the different screening strategies provide independent and complementary information, which can help to increase the overall number and diversity of the identified active compounds. Identifying structurally and chemically diverse active compounds with a similar binding mode is important for the creation of more comprehensive integrated pharmacophore models.

Limitations of the computational and experimental screening approach employed include a restricted scalability of the *in vitro* assay for assessing 3CLpro inhibitory activity, which has recently been addressed by the development of an optimized assay for high-throughput screening of 3CLpro [20], and limitations in terms of the size of the compound library and the conformational search space that could be explored using high-performance computing facilities available in a typical academic setting, which may be addressed using recent dedicated supercomputer-based screening approaches [24–26]. Extending 3CLpro virtual screening to supercomputer-based methodologies has also recently been shown to enable an improved modeling of receptor flexibility using ensemble docking [24], extensive MD simulations to generate high-resolution conformational ensembles [25], and the application of more advanced polarizable force fields [26].

As a next step, extended/further screening and structural optimization analyses for 3CLpro inhibitors will be required. This will help to pave the way for additional activity assessments in preclinical models and the subsequent development of lead compounds.

## Supporting information

SMILES strings for all experimentally confirmed active compounds

## DATA AND SOFTWARE AVAILABILITY

The Supporting Information contains the lists of all experimentally assessed candidate 3CLPro inhibitors, the dose-response curves and structures in SMILES format for all experimentally confirmed active compounds, the results for the validation analyses of the FRET-based 3CLPro *in vitro* assay and the reference inhibitor GC-376, and the detailed results for the physicochemical and ADME analyses. Scripts including the commands and parameters for the molecular docking analyses, the ligand similarity search, the machine learning based screening, the configuration settings used for the software NAMD, as well as a list of all used software tools and their versions, and exemplary video animations of MD simulations for confirmed inhibitors are available on GitHub (https://github.com/eglaab/3clpro).

## SUPPORTING INFORMATION

- Tables S1-S5: Lists of experimentally assessed candidate 3CLPro small molecule inhibitors
- Figures S1-S2: Validation of the FRET assay and the reference compound
- Figure S3: Dose-response curves for the active compounds
- Figures S4-S11: Physicochemical and ADME analyses for the active compounds
- Used software tools and versions for the virtual screening analyses
- Step-by-step description of the virtual screening analyses

## FUNDING SOURCES

Authors EG and DA received funding support from the Fonds National de la Recherche Luxembourg (FNR) as part of the COVID-19 Fast-Track research project CovScreen (COVID-19/2020-1/14715687).

## ACKNOWLEDGEMENTS

Bioinformatics analyses presented in this paper were carried out in part using the HPC facilities of the University of Luxembourg (see http://hpc.uni.lu).

## Notes

https://github.com/eglaab/3clpro

### Competing Interest Statement

The authors have declared no competing interest.

### Summary of Updates

Extensions and revisions have been included for the Introduction, Methods and Conclusions section, and the Supporting Information.

