## Supporting Information for "A Pharmacophore Model for SARS-CoV-2 3CLpro Small Molecule Inhibitors and in Vitro Experimental Validation of Computationally Screened Inhibitors"

### 1). Lists of experimentally assessed candidate 3CLPro inhibitors

**Table S1**: Previously reported small molecule inhibitors for 3CLPro for either SARS-CoV or SARS-CoV-2 derived from the literature and tested experimentally using a SARS-CoV-2 3CLPro activity assay. The compound name, the type of 3CLpro for which the compound was previously tested as inhibitor (Cov-1 = SARS-CoV, CoV-2 = SARS-CoV-2), the supplier compound ID (MolPort ID), the molecular weight (MW), the estimated n-octanol/water partition coefficient (derived from the ZINC database, or if not available, computed using the Molinspiration software, https://www.molinspiration.com), and the PubMed ID reference to the original publication in which the inhibition was reported (PMID). *Compounds highlighted by the star symbol have been confirmed as SARS-CoV-2 3CLpro inhibitors (see main manuscript).

| # | Compound name | 3CLPro SARS type | MolPort ID | MW | logP | Reference (PMID) |
|---|---|---|---|---|---|---|
| 1 | GC-376 / GC-363* | Cov-2 | 046-767-490 | 508 | -0.68 | 32887884 |
| 2 | Ebselen | Cov-2 | 003-666-383 | 274 | 2.92 | 32917717 |
| 3 | amentoflavone* | Cov-1 | 001-741-078 | 538 | 5.13 | 20934345 |
| 4 | hesperetine | Cov-1 | 001-741-418 | 302 | 2.52 | 16115693 |
| 5 | pectolinarin* | Cov-1 | 007-980-807 | 623 | -0.79 | 31724441 |
| 6 | baicalein* | Cov-2 | 001-741-408 | 270 | 2.58 | 32737471 |
| 7 | luteolin* | Cov-1 | 000-706-683 | 286 | 2.28 | 20934345 |
| 8 | quercetin | Cov-1 | 001-740-557 | 302 | 1.99 | 20934345 |
| 9 | pristimerin | Cov-1 | 001-740-445 | 465 | 6.79 | 20167482 |
| 10 | 1-Hydroxypyridine-2-thione zinc | Cov-1 | 023-220-327 | 318 | -0.46 | 15358550 |

**Table S2**: Candidate small molecule inhibitor compounds for SARS-CoV-2 3CLPro derived from the virtual screening of the ZINC and SWEETLEAD libraries (see Methods section in the main manuscript). The compound (IUPAC for uncharacterized compounds), the compound type (investigational drug, compound tested only in vivo, natural products, or uncharacterized; categories derived from the ZINC database), the supplier compound ID (MolPort ID), the molecular weight (MW), the estimated n-octanol/water partition coefficient (derived from the ZINC database, or if not available, computed using the Molinspiration software, https://www.molinspiration.com). *Compounds highlighted by the star symbol have been confirmed as SARS-CoV-2 3CLPro inhibitors (see main manuscript). Compounds displaying high background fluorescence in the 3CLpro activity assay were marked in blue in Tables S2 to S5.

| # | Compound name | Compound type | MolPort ID | MW | logP |
|---|---|---|---|---|---|
| 11 | Puerarin | investigational drug | 003-939-176 | 416 | 0.39 |
| 12 | Rottlerin* | in-vivo-only | 003-959-457 | 517 | 5.40 |
| 13 | 7-Hydroxy-8-[4-(2-hydroxy-ethyl)-piperazin-1-ylmethyl]-3-(4-methoxy-phenyl)-2-methyl-chromen-4-one | natural products | 002-360-471 | 424 | 2.59 |
| 14 | 3-(2-bromophenoxy)-8-{[4-(2-hydroxyethyl)piperazin-1-ium-1-yl]methyl}-2-methyl-4-oxo-4H-chromen-7-olate | uncharacterized | 000-773-665 | 489 | 3.47 |
| 15 | N-(9,10-dioxoanthryl)-2-[(3-hydroxypropyl)amino]acetamide | uncharacterized | 002-738-380 | 338 | 1.37 |
| 16 | 7-Hydroxy-8-[4-(2-hydroxy-ethyl)-piperazin-1-ylmethyl]-5-methoxy-2-phenyl-chromen-4-one* | natural products | 002-676-327 | 410 | 2.28 |
| 17 | N-(3-hydroxypropyl)-2-((4-methyl-6-oxo-6H-benzo[c]chromen-3-yl)oxy)propanamide | uncharacterized | 002-648-663 | 355 | 2.52 |
| 18 | N-(1-hydroxybutan-2-yl)-N'-(pyridin-3-yl)ethanediamide | uncharacterized | 003-648-158 | 237 | -0.09 |
| 19 | 1-{[3-(4-chlorophenyl)-7-oxido-2-oxochromen-8-yl]methyl}-4-(2-hydroxyethyl)piperazin-1-ium | uncharacterized | 000-842-586 | 415 | 2.93 |
| 20 | (2S,4S)-4-amino-N-methyl-1-[2-(4-methylbenzoyl)benzoyl]pyrrolidine-2-carboxamide | uncharacterized | 019-811-616 | 365 | 1.51 |
| 21 | 4-(2-{5,11-dioxo-5H,6H,11H-indeno[1,2-c]isoquinolin-6-yl}ethyl)benzene-1-sulfonamide | uncharacterized | 044-543-977 | 430 | 3.10 |
| 22 | N-[2-hydroxy-2-(4-hydroxyphenyl)ethyl]-N-methyl-2-[(4-oxo-3-phenyl-4H-chromen-7-yl)oxy]acetamide | uncharacterized | 000-848-761 | 445 | 3.74 |
| 23 | 4-{4,8-dioxo-3-phenyl-4H,8H,9H,10H-pyrano[2,3-h]chromen-10-yl}benzoic acid | uncharacterized | 039-055-985 | 412 | 4.60 |
| 24 | 3-(4-chlorophenyl)-2-[2-({6-oxo-6H-benzo[c]chromen-3-yl}oxy)acetamido]propanoic acid | uncharacterized | 002-525-292 | 452 | 3.79 |

| | | | | | |
|---|---|---|---|---|---|
| 25 | 4-[3-(4-hydroxyphenyl)-4,8-dioxo-4H,8H,9H,10H-pyrano[2,3-h]chromen-10-yl]benzoic acid | uncharacterized | 035-700-274 | 428 | 4.31 |
| 26 | 5-hydroxy-3-(2-methoxyphenyl)-10-(4-methoxyphenyl)-4H,8H,9H,10H-pyrano[2,3-h]chromene-4,8-dione | uncharacterized | 044-179-290 | 444 | 4.62 |
| 27 | 3-(4-chlorophenyl)-7-hydroxy-8-{[4-(2-hydroxyethyl)piperazin-1-yl]methyl}-4H-chromen-4-one | uncharacterized | 002-615-050 | 415 | 2.93 |
| 28 | N-methyl-2-({6-oxo-6H-benzo[c]chromen-3-yl}oxy)-N-(2,3,4,5,6-pentahydroxyhexyl)acetamide | uncharacterized | 002-536-294 | 447 | -0.78 |

**Table S3**: Candidate small molecule inhibitor compounds for SARS-CoV-2 3CLPro derived from the joint ligand-based and docking-based screening of the MolPort library (see Methods section in the main manuscript). The compound (IUPAC for uncharacterized compounds), the compound type (investigational drug, compound tested only in vivo, natural products, or uncharacterized; categories derived from the ZINC database), the supplier compound ID (MolPort ID), the molecular weight (MW), the estimated n-octanol/water partition coefficient (derived from the ZINC database, or if not available, computed using the Molinspiration software, https://www.molinspiration.com).

| # | Compound name | Compound type | MolPort ID | MW | logP |
|---|---|---|---|---|---|
| 29 | benzyl N-[(2S)-3-methyl-1-oxo-1-(propan-2-ylamino)butan-2-yl]carbamate | uncharacterized | 023-277-832 | 292 | 2.46 |
| 30 | benzyl N-[(1S)-2-methyl-1-(propylcarbamoyl)propyl]carbamate | uncharacterized | 023-277-830 | 292 | 2.46 |
| 31 | benzyl N-[(1S)-1-(butylcarbamoyl)-2-methylpropyl]carbamate | uncharacterized | 023-277-831 | 306 | 2.85 |
| 32 | (2S)-2-[(2S)-2-{[(benzyloxy)carbonyl]amino}-3-methylbutanamido]propanoic acid | uncharacterized | 028-959-397 | 322 | 1.53 |
| 33 | benzyl N-[(1S)-3-methyl-1-{[(1S)-3-methyl-1-{[(2S)-4-methyl-1-oxopentan-2-yl]carbamoyl}butyl]carbamoyl}butyl]carbamate | uncharacterized | 003-940-673 | 476 | 3.59 |

| | | | | | |
|---|---|---|---|---|---|
| 34 | 3-[(4-fluorophenyl)formamido]-N-[(3S)-2-oxoazepan-3-yl]propanamide | uncharacterized | 016-609-543 | 321 | 0.73 |
| 35 | N-[(2R)-2-hydroxypropyl]-3-methyl-2-(2-phenylacetamido)butanamide | uncharacterized | 030-011-861 | 292 | 0.87 |
| 36 | benzyl N-(1-{[(cyclopropylcarbamoyl)methyl](methyl)carbamoyl}-2-hydroxyethyl)carbamate | uncharacterized | 044-534-431 | 349 | 0.01 |
| 37 | N-[(2S)-2-hydroxypropyl]-2-(2-phenylacetamido)pentanamide | uncharacterized | 030-011-849 | 292 | 1.01 |
| 38 | N-[3-(2-oxopyrrolidin-1-yl)propyl]-3-(phenylformamido)butanamide | uncharacterized | 009-125-166 | 331 | 1.32 |
| 39 | 3-[1-(benzyloxy)propan-2-yl]-1-(2-oxoazepan-3-yl)urea | uncharacterized | 039-077-651 | 319 | 1.56 |
| 40 | N-(2-oxopiperidin-3-yl)-1-(2-phenylacetyl)piperidine-3-carboxamide | uncharacterized | 027-651-486 | 343 | 0.86 |
| 41 | N-[1-(oxolan-2-yl)ethyl]-2-(2-phenylacetamido)pentanamide | uncharacterized | 020-049-802 | 332 | 2.20 |
| 42 | benzyl N-(1-{[(tert-butylcarbamoyl)methyl]carbamoyl}-2-methylpropyl)carbamate | uncharacterized | 009-379-458 | 364 | 1.97 |
| 43 | benzyl (2S)-2-{[(5-oxopyrrolidin-3-yl)methyl]carbamoyl}pyrrolidine-1-carboxylate | uncharacterized | 042-601-620 | 345 | 1.04 |
| 44 | N-[2-(morpholin-4-yl)ethyl]-2-(2-phenylacetamido)pentanamide | uncharacterized | 006-419-396 | 348 | 0.96 |
| 45 | N-({[2-(morpholin-4-yl)ethyl]carbamoyl}methyl)-2-phenoxyacetamide | uncharacterized | 003-250-889 | 321 | -0.37 |
| 46 | benzyl N-[1-(cyclopropylcarbamoyl)-2-methylbutyl]carbamate | uncharacterized | 005-601-702 | 304 | 2.61 |
| 47 | 2-[2-(2-phenoxyacetamido)acetamido]-N-propylpropanamide | uncharacterized | 005-626-882 | 321 | 0.21 |
| 48 | N-[(cyclopropylcarbamoyl)methyl]-2-[(2,4-dichlorophenyl)formamido]-3-methylbutanamide | uncharacterized | 005-602-029 | 386 | 2.14 |
| 49 | N-(1-{4-[(cyclopropylcarbamoyl)methyl]pi | uncharacterized | 004-242-605 | 401 | 0.79 |

| | | | | | |
|---|---|---|---|---|---|
| | perazin-1-yl}-3-methyl-1-oxobutan-2-yl)-2-phenylacetamide | | | | |
| 50 | N-[2-(oxolan-2-yl)ethyl]-2-(2-phenylacetamido)pentanamide | uncharacterized | 020-049-873 | 332 | 2.20 |
| 51 | 1-[(benzylcarbamoyl)amino]-N-(2-oxoazepan-3-yl)cyclopentane-1-carboxamide | uncharacterized | 027-690-491 | 373 | 1.58 |
| 52 | benzyl N-[(1S)-1-[({hexahydro-1H-pyrrolo[2,1-c][1,4]oxazin-3-yl}methyl)carbamoyl]-2-methylpropyl]carbamate | uncharacterized | 009-113-309 | 390 | 1.92 |
| 53 | N-cyclopropyl-2-{[1-(3-phenoxypropanoyl)piperidin-3-yl]formamido}acetamide | uncharacterized | 009-093-573 | 374 | 1.92 |
| 54 | benzyl N-[(1S)-3-methyl-1-{[(1R)-3-methyl-1-{[(2S)-4-methyl-1-oxopentan-2-yl]carbamoyl}butyl]carbamoyl}butyl]carbamate | uncharacterized | 009-019-420 | 476 | 3.59 |
| 55 | 2-[(2,4-dichlorophenyl)formamido]-N-[1-(ethylcarbamoyl)ethyl]-4-methylpentanamide | uncharacterized | 005-574-787 | 402 | 2.78 |
| 56 | N-({[3-(morpholin-4-yl)propyl]carbamoyl}methyl)-2-phenoxyacetamide | uncharacterized | 003-251-113 | 335 | 0.02 |
| 57 | benzyl N-[2-methyl-1-(propylcarbamoyl)propyl]carbamate | uncharacterized | 005-669-468 | 292 | 2.46 |
| 58 | benzyl N-[1-(cyclopropylcarbamoyl)-2-methylpropyl]carbamate | uncharacterized | 005-597-829 | 290 | 2.22 |

**Table S4**: Candidate small molecule inhibitor compounds for SARS-CoV-2 3CLPro derived from the screening of the MolPort library based purely on molecular docking (see Methods section in the main manuscript). The compound (IUPAC for uncharacterized compounds), the compound type (investigational drug, compound tested only in vivo, natural products, or uncharacterized; categories derived from the ZINC database), the supplier compound ID (MolPort ID), the molecular weight (MW), the estimated n-octanol/water partition coefficient (derived from the ZINC database, or if not available, computed using the Molinspiration software, https://www.molinspiration.com). *Compounds highlighted by the star symbol have been confirmed as SARS-CoV-2 3CLPro inhibitors (see main manuscript).

| # | Compound name | Compound type | MolPort ID | MW | logP |
|---|---|---|---|---|---|
| 59 | N-{[1-(4-chlorophenyl)-7,7-dimethyl-2,5-dioxo-1,2,5,6,7,8-hexahydroquinolin-3-yl]formamido}-2-methylanilinium chloride | uncharacterized | 000-699-391 | 486 | 4.71 |
| 60 | 2-[4-(2-hydroxyethyl)piperazine-1-carbonyl]-3H-benzo[f]chromen-3-one | uncharacterized | 000-556-234 | 352 | 1.70 |
| 61 | 2-[(2-{[(9-ethyl-9H-carbazol-3-yl)methyl]amino}ethyl)amino]ethan-1-ol | uncharacterized | 000-124-705 | 311 | 2.49 |
| 62 | N-(2-benzoylphenyl)-2-(pyrrolidin-1-yl)acetamide | uncharacterized | 000-384-806 | 308 | 2.95 |
| 63 | 1-{[3-(4-bromophenyl)-7-oxido-4-oxo-4H-chromen-8-yl]methyl}-4-(2-hydroxyethyl)piperazin-1-ium | uncharacterized | 000-778-254 | 459 | 3.04 |
| 64 | 1-{[3-(4-chlorophenyl)-7-oxido-2-oxo-2H-chromen-8-yl]methyl}-4-(2-hydroxyethyl)piperazin-1-ium | uncharacterized | 000-842-586 | 415 | 2.93 |
| 65 | N-(2-benzoylphenyl)-2-(morpholin-4-yl)acetamide | uncharacterized | 000-384-709 | 324 | 2.19 |
| 66 | N-(5-{[(naphthalen-1-yl)methyl]sulfanyl}-1H-1,2,4-triazol-3-yl)acetamide | uncharacterized | 000-799-813 | 298 | 3.86 |
| 67 | ethyl 8-methyl-4-{[4-(2-methylpropoxy)phenyl]amino}quinoline-3-carboxylate | uncharacterized | 000-665-928 | 379 | 5.50 |
| 68 | N-phenyl-2-(piperazin-1-yl)acetamide dihydrochloride | uncharacterized | 000-158-251 | 256 | 0.53 |
| 69 | N-(2-aminoethyl)-2-{6-ethyl-12,12-dimethyl-3,5-dioxo-11-oxa-8-thia-4,6-diazatricyclo[7.4.0.0$^{2}$,$^{7}$]trideca-1(9),2(7)-dien-4-yl}acetamide | uncharacterized | 000-849-001 | 381 | 0.17 |
| 70 | 2-methyl-2-[({5-[3-(trifluoromethyl)phenyl]furan-2-yl}methyl)amino]propan-1-ol | uncharacterized | 000-137-291 | 313 | 3.83 |
| 71 | 2-{[(9-ethyl-9H-carbazol-3-yl)methyl]amino}ethan-1-ol | uncharacterized | 000-124-698 | 305 | 2.81 |
| 72 | 3-[(5E)-5-[(3,4-dihydroxyphenyl)methylidene]-4-oxo-2-sulfanylidene-1,3-thiazolidin-3-yl]benzoic acid* | uncharacterized | 000-821-805 | 373 | 3.20 |

| | | | | | |
|---|---|---|---|---|---|
| 73 | 3-amino-1-(naphthalen-1-yl)thiourea | uncharacterized | 000-157-715 | 217 | 2.00 |
| 74 | 4-(2-hydroxyethyl)-1-{[3-(4-methoxyphenyl)-7-oxido-2-oxo-2H-chromen-8-yl]methyl}piperazin-1-ium | uncharacterized | 000-844-215 | 411 | 2.28 |
| 75 | 1-(2,3,4,9-tetrahydro-1H-carbazol-6-yl)methanamine | uncharacterized | 000-226-151 | 200 | 2.51 |
| 76 | 3-amino-1-[4-(trifluoromethyl)phenyl]thiourea | uncharacterized | 000-159-120 | 235 | 1.87 |

**Table S5**: Candidate small molecule inhibitor compounds for SARS-CoV-2 3CLPro derived from the screening of the MolPort library based purely on machine learning (see Methods section in the main manuscript). The compound (IUPAC for uncharacterized compounds), the compound type (investigational drug, compound tested only in vivo, natural products, or uncharacterized; categories derived from the ZINC database), the supplier compound ID (MolPort ID), the molecular weight (MW), the estimated n-octanol/water partition coefficient (derived from the ZINC database, or if not available, computed using the Molinspiration software, https://www.molinspiration.com). *Compounds highlighted by the star symbol have been confirmed as SARS-CoV-2 3CLPro inhibitors (see main manuscript).

| # | Compound name | Compound type | MolPort ID | MW | logP |
|---|---|---|---|---|---|
| 77 | (2R)-N-tert-butyl-2-[N-(4-tert-butylphenyl)-1-(furan-2-yl)formamido]-2-(pyridin-3-yl)acetamide* | uncharacterized | 047-408-524 | 434 | 4.67 |
| 78 | ethyl (2E,4S)-4-[(2R,5S)-2-[(4-fluorophenyl)methyl]-6-methyl-5-[(5-methyl-1,2-oxazol-3-yl)formamido]-4-oxoheptanamido]-5-[(3S)-2-oxopyrrolidin-3-yl]pent-2-enoate | uncharacterized | 006-170-056 | 599 | 2.83 |
| 79 | N-(3-acetamidophenyl)-3-(1H-indol-3-yl)-2-[(thiophen-2-yl)formamido]propanamide | uncharacterized | 005-779-727 | 447 | 4.17 |
| 80 | 4-[3-(1H-indol-3-yl)-2-[(thiophen-2-yl)formamido]propanamido]-N-methylbenzamide | uncharacterized | 004-604-653 | 447 | 3.57 |
| 81 | 3-[3-(1H-indol-3-yl)-2-[(thiophen-2-yl)formamido]propanamido]-N-methylbenzamide | uncharacterized | 005-769-464 | 447 | 3.57 |

| | | | | | |
|---|---|---|---|---|---|
| 82 | (2S)-3-(1H-indol-3-yl)-N-[4-(2-oxopyrrolidin-1-yl)phenyl]-2-[(thiophen-2-yl)formamido]propanamide | uncharacterized | 005-785-350 | 473 | 4.34 |
| 83 | N-[(1S)-1-(1H-1,3-benzodiazol-2-yl)-2-phenylethyl]-2-{[4-(2-oxopyrrolidin-1-yl)phenyl]formamido}acetamide | uncharacterized | 007-308-518 | 482 | 3.52 |
| 84 | ethyl 3-(3-{3-[(thiophen-2-yl)methyl]-3H-imidazo[4,5-b]pyridin-2-yl}propanamido)benzoate | uncharacterized | 007-650-576 | 435 | 4.29 |
| 85 | N-[1-(1H-1,3-benzodiazol-2-yl)-2-phenylethyl]-2-{[4-(2-oxopyrrolidin-1-yl)phenyl]formamido}acetamide | uncharacterized | 009-372-692 | 482 | 3.52 |
| 86 | 2-[2-(1H-1,2,3-benzotriazol-1-yl)-N-benzylacetamido]-N-(4-methoxyphenyl)-2-(thiophen-2-yl)acetamide | uncharacterized | 003-851-907 | 512 | 4.91 |
| 87 | N-cyclopropyl-3-[3-(1H-indol-3-yl)-2-[(thiophen-2-yl)formamido]propanamido]benzamide | uncharacterized | 004-562-271 | 473 | 4.10 |
| 88 | N,N-diethyl-3-[3-(1H-indol-3-yl)-2-[(thiophen-2-yl)formamido]propanamido]benzamide | uncharacterized | 004-591-387 | 489 | 4.69 |
| 89 | ethyl 4-(3-{3-[(thiophen-2-yl)methyl]-3H-imidazo[4,5-b]pyridin-2-yl}propanamido)benzoate | uncharacterized | 007-650-575 | 435 | 4.29 |
| 90 | (4R,7R,10R)-10-benzyl-4-methyl-15-[3-(1-methyl-1H-pyrazol-5-yl)propanoyl]-7-(2-methylpropyl)-18-oxa-3,6,9,12,15,19-hexaazabicyclo[15.2.1]icosa-1(19),17(20)-diene-2,5,8,11-tetrone | uncharacterized | 046-756-573 | 635 | 0.88 |
| 91 | 3-(1H-indol-3-yl)-N-{3-[3-(2-methylpropyl)-1,2,4-oxadiazol-5-yl]propyl}-2-[(thiophen-2-yl)formamido]propanamide | uncharacterized | 009-107-411 | 480 | 3.90 |

| | | | | | |
|---|---|---|---|---|---|
| 92 | N-{1-[1-({[(thiophen-2-yl)methyl]carbamoyl}methyl)-1H-1,3-benzodiazol-2-yl]butyl}benzamide | uncharacterized | 028-810-678 | 447 | 4.69 |
| 93 | (4S,7S,10R)-7,10-dibenzyl-15-cyclobutanecarbonyl-4-methyl-18-oxa-3,6,9,12,15,19-hexaazabicyclo[15.2.1]icosa-1(19),17(20)-diene-2,5,8,11-tetrone | uncharacterized | 046-753-433 | 615 | 1.51 |
| 94 | 2-[2-(1H-1,2,3-benzotriazol-1-yl)-N-[(furan-2-yl)methyl]acetamido]-N-benzyl-2-(3-methoxyphenyl)acetamide | uncharacterized | 001-997-526 | 510 | 4.12 |
| 95 | 3-(1H-indol-3-yl)-N-[3-(2-oxopyrrolidin-1-yl)phenyl]-2-[(thiophen-2-yl)formamido]propanamide | uncharacterized | 009-379-497 | 473 | 4.34 |
| 96 | 2-[2-(1H-1,2,3-benzotriazol-1-yl)-N-[(furan-2-yl)methyl]acetamido]-N-benzyl-2-(4-methylphenyl)acetamide | uncharacterized | 001-997-520 | 494 | 4.42 |

### 3.) Validation of the assay and the reference compound

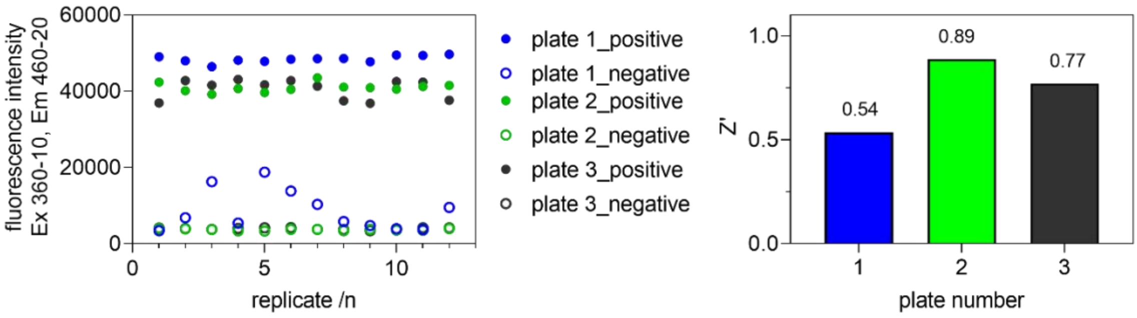


Figure S1: Validation of the established FRET-based 3CLPro *in vitro* assay. The assay displayed a Z’ value of > 0.5 and is suitable for the assessment of inhibitors with sub µM to low mM $IC_{50}$ values. ‘Positive’ indicates the fluorescence intensity of the complex between 3CLpro and the substrate peptide, whereas ‘negative’ indicates the background fluorescence resulting from the substrate peptide.

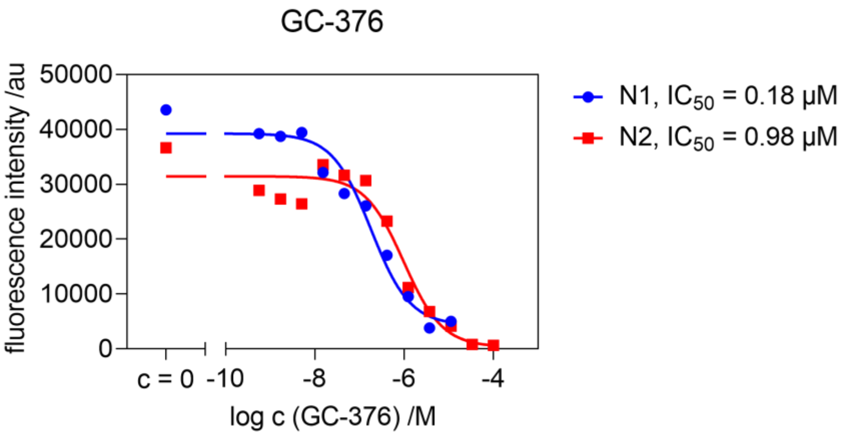


Figure S2: Validation of the reference compound GC-376. The $IC_{50}$ of GC-376 is estimated to be in the range of 0.18 to 0.98 µM, in line with the previously reported $IC_{50}$ values for GC-376 (0.19 µM) and GC-373 (0.4 µM) in the literature (Vuong et al., Nat. Comm., 2020, doi: 10.1038/s41467-020-18096-2).

#### 4.) Dose-response curves

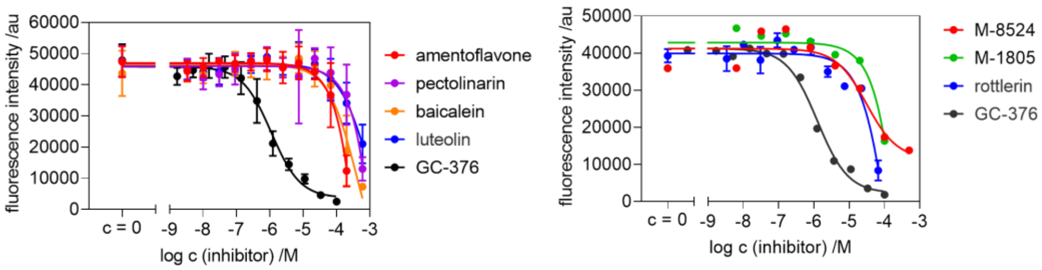


Figure S3: Dose-response curves obtained from the FRET-based 3CLpro *in vitro* assay for the selected compounds.

#### 5.) Physicochemical and ADME analyses

Physicochemical and ADME characteristics of the confirmed small molecule inhibitors for 3CLPro, derived using the software SwissADME:

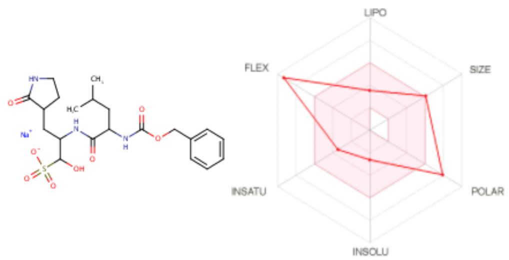


SMILES CC(CC(C(=O)NC(C(S(=O)(=O)[O-])O)CC1CCNC1=O)NC(=O)OCc1ccccc1)C.[Na+]

| Physicochemical Properties | |
|---|---|
| Formula | C21H30N3NaO8S |
| Molecular weight | 507.53 g/mol |
| Num. heavy atoms | 34 |
| Num. arom. heavy atoms | 6 |
| Fraction Csp3 | 0.57 |
| Num. rotatable bonds | 14 |
| Num. H-bond acceptors | 8 |
| Num. H-bond donors | 4 |
| Molar Refractivity | 120.83 |
| TPSA | 182.34 Å² |

| Lipophilicity | |
|---|---|
| Log $P_{o/w}$ (iLOGP) | -12.77 |
| Log $P_{o/w}$ (XLOGP3) | 0.71 |
| Log $P_{o/w}$ (WLOGP) | 0.75 |
| Log $P_{o/w}$ (MLOGP) | -0.08 |
| Log $P_{o/w}$ (SILICOS-IT) | 0.44 |
| Consensus Log $P_{o/w}$ | -2.19 |

| Water Solubility | |
|---|---|
| Log *S* (ESOL) | -2.64 |
| Solubility | 1.16e+00 mg/ml ; 2.29e-03 mol/l |
| Class | Soluble |
| Log *S* (Ali) | -4.12 |
| Solubility | 3.88e-02 mg/ml ; 7.64e-05 mol/l |
| Class | Moderately soluble |
| Log *S* (SILICOS-IT) | -3.75 |
| Solubility | 8.94e-02 mg/ml ; 1.76e-04 mol/l |
| Class | Soluble |

| Pharmacokinetics | |
|---|---|
| GI absorption | Low |
| BBB permeant | No |
| P-gp substrate | Yes |
| CYP1A2 inhibitor | No |
| CYP2C19 inhibitor | No |
| CYP2C9 inhibitor | No |
| CYP2D6 inhibitor | No |
| CYP3A4 inhibitor | No |
| Log $K_p$ (skin permeation) | -8.89 cm/s |

| Druglikeness | |
|---|---|
| Lipinski | No; 2 violations: MW>500, NorO>10 |
| Ghose | No; 1 violation: MW>480 |
| Veber | No; 2 violations: Rotors>10, TPSA>140 |
| Egan | No; 1 violation: TPSA>131.6 |
| Muegge | No; 1 violation: TPSA>150 |
| Bioavailability Score | 0.17 |

| Medicinal Chemistry | |
|---|---|
| PAINS | 0 alert |
| Brenk | 1 alert: sulfonic_acid_2 |
| Leadlikeness | No; 2 violations: MW>350, Rotors>7 |
| Synthetic accessibility | 5.01 |

Figure S4: Physicochemical and ADME properties for compound GC-376

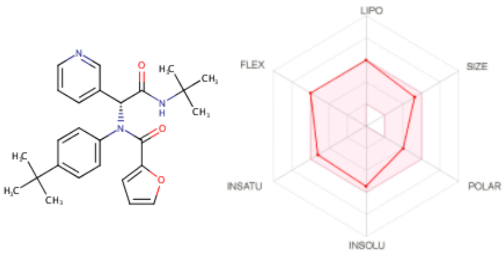


SMILES O=C(N([C@@H](C(=O)NC(C)(C)C)c1cccnc1)c1ccc(cc1)C(C)(C)C)c1ccco1

| Physicochemical Properties | |
|---|---|
| Formula | C26H31N3O3 |
| Molecular weight | 433.54 g/mol |
| Num. heavy atoms | 32 |
| Num. arom. heavy atoms | 17 |
| Fraction Csp3 | 0.35 |
| Num. rotatable bonds | 9 |
| Num. H-bond acceptors | 4 |
| Num. H-bond donors | 1 |
| Molar Refractivity | 126.45 |
| TPSA | 75.44 Å² |

| Lipophilicity | |
|---|---|
| Log $P_{o/w}$ (iLOGP) | 4.21 |
| Log $P_{o/w}$ (XLOGP3) | 4.97 |
| Log $P_{o/w}$ (WLOGP) | 4.95 |
| Log $P_{o/w}$ (MLOGP) | 2.38 |
| Log $P_{o/w}$ (SILICOS-IT) | 4.44 |
| Consensus Log $P_{o/w}$ | 4.19 |

| Water Solubility | |
|---|---|
| Log *S* (ESOL) | -5.46 |
| Solubility | 1.51e-03 mg/ml ; 3.48e-06 mol/l |
| Class | Moderately soluble |
| Log *S* (Ali) | -6.29 |
| Solubility | 2.21e-04 mg/ml ; 5.10e-07 mol/l |
| Class | Poorly soluble |
| Log *S* (SILICOS-IT) | -7.92 |
| Solubility | 5.24e-06 mg/ml ; 1.21e-08 mol/l |
| Class | Poorly soluble |

| Pharmacokinetics | |
|---|---|
| GI absorption | High |
| BBB permeant | No |
| P-gp substrate | No |
| CYP1A2 inhibitor | No |
| CYP2C19 inhibitor | Yes |
| CYP2C9 inhibitor | Yes |
| CYP2D6 inhibitor | Yes |
| CYP3A4 inhibitor | Yes |
| Log $K_p$ (skin permeation) | -5.42 cm/s |

| Druglikeness | |
|---|---|
| Lipinski | Yes; 0 violation |
| Ghose | Yes |
| Veber | Yes |
| Egan | Yes |
| Muegge | Yes |
| Bioavailability Score | 0.55 |

| Medicinal Chemistry | |
|---|---|
| PAINS | 0 alert |
| Brenk | 0 alert |
| Leadlikeness | No; 3 violations: MW>350, Rotors>7, XLOGP3>3.5 |
| Synthetic accessibility | 3.92 |

Figure S5: Physicochemical and ADME properties for compound M-8524 (MolPort ID: 047-408-524)

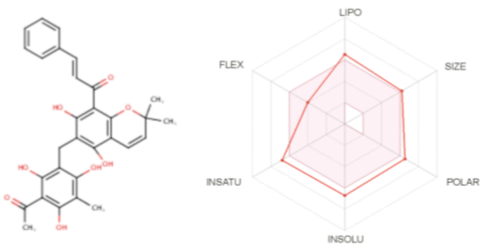


SMILES O=C(c1c(O)c(Cc2c(O)c(C)c(c(c2O)C(=O)C)O)c(c2c1OC(C)(C)C=C2)O)/C=C/c1ccccc1

| Physicochemical Properties | |
|---|---|
| Formula | C30H28O8 |
| Molecular weight | 516.54 g/mol |
| Num. heavy atoms | 38 |
| Num. arom. heavy atoms | 18 |
| Fraction Csp3 | 0.20 |
| Num. rotatable bonds | 6 |
| Num. H-bond acceptors | 8 |
| Num. H-bond donors | 5 |
| Molar Refractivity | 145.10 |
| TPSA | 144.52 Å² |

| Lipophilicity | |
|---|---|
| Log $P_{o/w}$ (iLOGP) | 3.76 |
| Log $P_{o/w}$ (XLOGP3) | 5.91 |
| Log $P_{o/w}$ (WLOGP) | 5.18 |
| Log $P_{o/w}$ (MLOGP) | 1.66 |
| Log $P_{o/w}$ (SILICOS-IT) | 5.55 |
| Consensus Log $P_{o/w}$ | 4.41 |

| Water Solubility | |
|---|---|
| Log *S* (ESOL) | -6.72 |
| Solubility | 9.83e-05 mg/ml ; 1.90e-07 mol/l |
| Class | Poorly soluble |
| Log *S* (Ali) | -8.72 |
| Solubility | 9.87e-07 mg/ml ; 1.91e-09 mol/l |
| Class | Poorly soluble |
| Log *S* (SILICOS-IT) | -6.39 |
| Solubility | 2.09e-04 mg/ml ; 4.06e-07 mol/l |
| Class | Poorly soluble |

| Pharmacokinetics | |
|---|---|
| GI absorption | Low |
| BBB permeant | No |
| P-gp substrate | No |
| CYP1A2 inhibitor | No |
| CYP2C19 inhibitor | Yes |
| CYP2C9 inhibitor | No |
| CYP2D6 inhibitor | No |
| CYP3A4 inhibitor | No |
| Log $K_p$ (skin permeation) | -5.25 cm/s |

| Druglikeness | |
|---|---|
| Lipinski | Yes; 1 violation: MW>500 |
| Ghose | No; 2 violations: MW>480, MR>130 |
| Veber | No; 1 violation: TPSA>140 |
| Egan | No; 1 violation: TPSA>131.6 |
| Muegge | No; 1 violation: XLOGP3>5 |
| Bioavailability Score | 0.55 |

| Medicinal Chemistry | |
|---|---|
| PAINS | 0 alert |
| Brenk | 1 alert: michael_acceptor_1 |
| Leadlikeness | No; 2 violations: MW>350, XLOGP3>3.5 |
| Synthetic accessibility | 4.57 |

Figure S6: Physicochemical and ADME properties for compound rottlerin

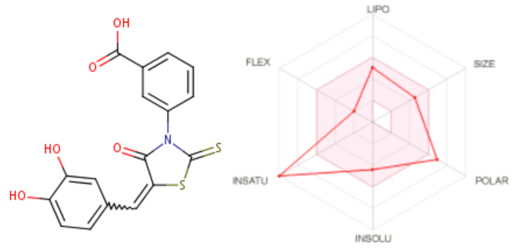


SMILES OC(=O)c1cccc(c1)N1C(=S)S/C(=C/c2ccc(c(c2)O)O)/C1=O

| Physicochemical Properties | |
|---|---|
| Formula | C17H11NO5S2 |
| Molecular weight | 373.40 g/mol |
| Num. heavy atoms | 25 |
| Num. arom. heavy atoms | 12 |
| Fraction Csp3 | 0.00 |
| Num. rotatable bonds | 3 |
| Num. H-bond acceptors | 5 |
| Num. H-bond donors | 3 |
| Molar Refractivity | 102.48 |
| TPSA | 155.46 Å² |

| Lipophilicity | |
|---|---|
| Log $P_{o/w}$ (iLOGP) | 1.90 |
| Log $P_{o/w}$ (XLOGP3) | 3.35 |
| Log $P_{o/w}$ (WLOGP) | 2.71 |
| Log $P_{o/w}$ (MLOGP) | 1.38 |
| Log $P_{o/w}$ (SILICOS-IT) | 3.10 |
| Consensus Log $P_{o/w}$ | 2.49 |

| Water Solubility | |
|---|---|
| Log *S* (ESOL) | -4.42 |
| Solubility | 1.41e-02 mg/ml ; 3.78e-05 mol/l |
| Class | Moderately soluble |
| Log *S* (Ali) | -6.29 |
| Solubility | 1.91e-04 mg/ml ; 5.10e-07 mol/l |
| Class | Poorly soluble |
| Log *S* (SILICOS-IT) | -3.37 |
| Solubility | 1.59e-01 mg/ml ; 4.27e-04 mol/l |
| Class | Soluble |

| Pharmacokinetics | |
|---|---|
| GI absorption | Low |
| BBB permeant | No |
| P-gp substrate | No |
| CYP1A2 inhibitor | Yes |
| CYP2C19 inhibitor | No |
| CYP2C9 inhibitor | Yes |
| CYP2D6 inhibitor | No |
| CYP3A4 inhibitor | Yes |
| Log $K_p$ (skin permeation) | -6.20 cm/s |

| Druglikeness | |
|---|---|
| Lipinski | Yes; 0 violation |
| Ghose | Yes |
| Veber | No; 1 violation: TPSA>140 |
| Egan | No; 1 violation: TPSA>131.6 |
| Muegge | No; 1 violation: TPSA>150 |
| Bioavailability Score | 0.11 |

| Medicinal Chemistry | |
|---|---|
| PAINS | 2 alerts: catechol_A, ene_rhod_A |
| Brenk | 3 alerts: catechol, michael_acceptor_1, thiocarbonyl_group |
| Leadlikeness | No; 1 violation: MW>350 |
| Synthetic accessibility | 3.16 |

Figure S7: Physicochemical and ADME properties for compound M-1805 (MolPort ID: 000-821-805)

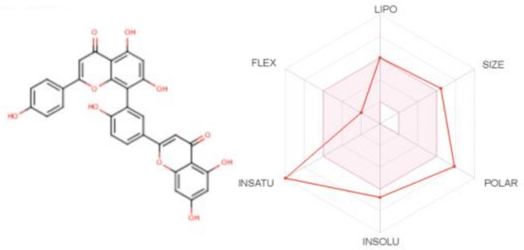


SMILES Oc1ccc(cc1)c1cc(=O)c2c(o1)c(c(cc2O)O)c1cc(ccc1O)c1cc(=O)c2c(o1)cc(cc2O)O

| Physicochemical Properties | |
|---|---|
| Formula | C30H18O10 |
| Molecular weight | 538.46 g/mol |
| Num. heavy atoms | 40 |
| Num. arom. heavy atoms | 32 |
| Fraction Csp3 | 0.00 |
| Num. rotatable bonds | 3 |
| Num. H-bond acceptors | 10 |
| Num. H-bond donors | 6 |
| Molar Refractivity | 146.97 |
| TPSA | 181.80 Å² |

| Lipophilicity | |
|---|---|
| Log $P_{o/w}$ (iLOGP) | 3.06 |
| Log $P_{o/w}$ (XLOGP3) | 5.04 |
| Log $P_{o/w}$ (WLOGP) | 5.13 |
| Log $P_{o/w}$ (MLOGP) | 0.25 |
| Log $P_{o/w}$ (SILICOS-IT) | 4.61 |
| Consensus Log $P_{o/w}$ | 3.62 |

| Water Solubility | |
|---|---|
| Log *S* (ESOL) | -6.75 |
| Solubility | 9.63e-05 mg/ml ; 1.79e-07 mol/l |
| Class | Poorly soluble |
| Log *S* (Ali) | -8.60 |
| Solubility | 1.36e-06 mg/ml ; 2.52e-09 mol/l |
| Class | Poorly soluble |
| Log *S* (SILICOS-IT) | -8.70 |
| Solubility | 1.07e-06 mg/ml ; 1.98e-09 mol/l |
| Class | Poorly soluble |

| Pharmacokinetics | |
|---|---|
| GI absorption | Low |
| BBB permeant | No |
| P-gp substrate | No |
| CYP1A2 inhibitor | No |
| CYP2C19 inhibitor | No |
| CYP2C9 inhibitor | No |
| CYP2D6 inhibitor | No |
| CYP3A4 inhibitor | No |
| Log $K_p$ (skin permeation) | -6.01 cm/s |

| Druglikeness | |
|---|---|
| Lipinski | No; 2 violations: MW>500, NHorOH>5 |
| Ghose | No; 2 violations: MW>480, MR>130 |
| Veber | No; 1 violation: TPSA>140 |
| Egan | No; 1 violation: TPSA>131.6 |
| Muegge | No; 3 violations: XLOGP3>5, TPSA>150, H-don>5 |
| Bioavailability Score | 0.17 |

| Medicinal Chemistry | |
|---|---|
| PAINS | 0 alert |
| Brenk | 0 alert |
| Leadlikeness | No; 2 violations: MW>350, XLOGP3>3.5 |
| Synthetic accessibility | 4.27 |

Figure S8: Physicochemical and ADME properties for compound amentoflavone

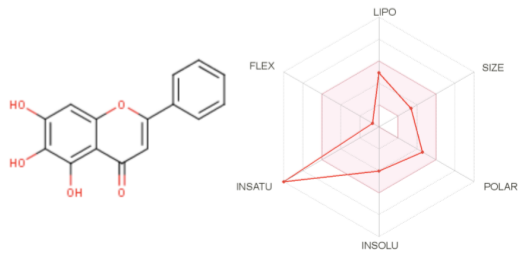


SMILES Oc1cc2oc(cc(=O)c2c(c1O)O)c1ccccc1

| Physicochemical Properties | |
|---|---|
| Formula | C15H10O5 |
| Molecular weight | 270.24 g/mol |
| Num. heavy atoms | 20 |
| Num. arom. heavy atoms | 16 |
| Fraction Csp3 | 0.00 |
| Num. rotatable bonds | 1 |
| Num. H-bond acceptors | 5 |
| Num. H-bond donors | 3 |
| Molar Refractivity | 73.99 |
| TPSA | 90.90 Å² |

| Lipophilicity | |
|---|---|
| Log $P_{o/w}$ (iLOGP) | 2.43 |
| Log $P_{o/w}$ (XLOGP3) | 3.16 |
| Log $P_{o/w}$ (WLOGP) | 2.58 |
| Log $P_{o/w}$ (MLOGP) | 0.52 |
| Log $P_{o/w}$ (SILICOS-IT) | 2.52 |
| Consensus Log $P_{o/w}$ | 2.24 |

| Water Solubility | |
|---|---|
| Log *S* (ESOL) | -4.03 |
| Solubility | 2.51e-02 mg/ml ; 9.28e-05 mol/l |
| Class | Moderately soluble |
| Log *S* (Ali) | -4.74 |
| Solubility | 4.93e-03 mg/ml ; 1.82e-05 mol/l |
| Class | Moderately soluble |
| Log *S* (SILICOS-IT) | -4.40 |
| Solubility | 1.07e-02 mg/ml ; 3.94e-05 mol/l |
| Class | Moderately soluble |

| Pharmacokinetics | |
|---|---|
| GI absorption | High |
| BBB permeant | No |
| P-gp substrate | No |
| CYP1A2 inhibitor | Yes |
| CYP2C19 inhibitor | No |
| CYP2C9 inhibitor | No |
| CYP2D6 inhibitor | Yes |
| CYP3A4 inhibitor | Yes |
| Log $K_p$ (skin permeation) | -5.70 cm/s |

| Druglikeness | |
|---|---|
| Lipinski | Yes; 0 violation |
| Ghose | Yes |
| Veber | Yes |
| Egan | Yes |
| Muegge | Yes |
| Bioavailability Score | 0.55 |

| Medicinal Chemistry | |
|---|---|
| PAINS | 1 alert: catechol_A |
| Brenk | 1 alert: catechol |
| Leadlikeness | Yes |
| Synthetic accessibility | 3.02 |

Figure S9: Physicochemical and ADME properties for compound baicalein

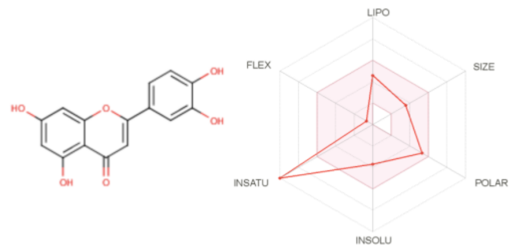


SMILES Oc1cc(O)c2c(c1)oc(cc2=O)c1ccc(c(c1)O)O

| Physicochemical Properties | |
|---|---|
| Formula | C15H10O6 |
| Molecular weight | 286.24 g/mol |
| Num. heavy atoms | 21 |
| Num. arom. heavy atoms | 16 |
| Fraction Csp3 | 0.00 |
| Num. rotatable bonds | 1 |
| Num. H-bond acceptors | 6 |
| Num. H-bond donors | 4 |
| Molar Refractivity | 76.01 |
| TPSA | 111.13 Å² |

| Lipophilicity | |
|---|---|
| Log $P_{o/w}$ (iLOGP) | 1.86 |
| Log $P_{o/w}$ (XLOGP3) | 2.53 |
| Log $P_{o/w}$ (WLOGP) | 2.28 |
| Log $P_{o/w}$ (MLOGP) | -0.03 |
| Log $P_{o/w}$ (SILICOS-IT) | 2.03 |
| Consensus Log $P_{o/w}$ | 1.73 |

| Water Solubility | |
|---|---|
| Log *S* (ESOL) | -3.71 |
| Solubility | 5.63e-02 mg/ml ; 1.97e-04 mol/l |
| Class | Soluble |
| Log *S* (Ali) | -4.51 |
| Solubility | 8.84e-03 mg/ml ; 3.09e-05 mol/l |
| Class | Moderately soluble |
| Log *S* (SILICOS-IT) | -3.82 |
| Solubility | 4.29e-02 mg/ml ; 1.50e-04 mol/l |
| Class | Soluble |

| Pharmacokinetics | |
|---|---|
| GI absorption | High |
| BBB permeant | No |
| P-gp substrate | No |
| CYP1A2 inhibitor | Yes |
| CYP2C19 inhibitor | No |
| CYP2C9 inhibitor | No |
| CYP2D6 inhibitor | Yes |
| CYP3A4 inhibitor | Yes |
| Log $K_p$ (skin permeation) | -6.25 cm/s |

| Druglikeness | |
|---|---|
| Lipinski | Yes; 0 violation |
| Ghose | Yes |
| Veber | Yes |
| Egan | Yes |
| Muegge | Yes |
| Bioavailability Score | 0.55 |

| Medicinal Chemistry | |
|---|---|
| PAINS | 1 alert: catechol_A |
| Brenk | 1 alert: catechol |
| Leadlikeness | Yes |
| Synthetic accessibility | 3.02 |

Figure S10: Physicochemical and ADME properties for compound luteolin

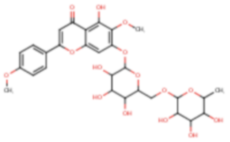

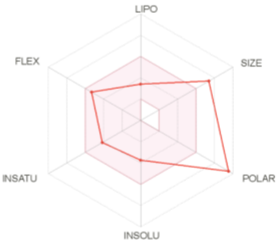


| | |
|---|---|
| SMILES | COc1c(OC2OC(COC3OC(C)C(C(C3O)O)O)C(C(C2O)O)O)cc2c(c1O)c(=O)cc(o2)c1ccc(cc1)OC |
| **Physicochemical Properties** | |
| Formula | C29H34O15 |
| Molecular weight | 622.57 g/mol |
| Num. heavy atoms | 44 |
| Num. arom. heavy atoms | 16 |
| Fraction Csp3 | 0.48 |
| Num. rotatable bonds | 8 |
| Num. H-bond acceptors | 15 |
| Num. H-bond donors | 7 |
| Molar Refractivity | 148.29 |
| TPSA | 227.20 Å² |
| **Lipophilicity** | |
| Log $P_{o/w}$ (iLOGP) | 1.73 |
| Log $P_{o/w}$ (XLOGP3) | 0.47 |
| Log $P_{o/w}$ (WLOGP) | -0.79 |
| Log $P_{o/w}$ (MLOGP) | -3.03 |
| Log $P_{o/w}$ (SILICOS-IT) | -0.53 |
| Consensus Log $P_{o/w}$ | -0.43 |

| | |
|---|---|
| **Water Solubility** | |
| Log *S* (ESOL) | -3.74 |
| Solubility | 1.14e-01 mg/ml ; 1.83e-04 mol/l |
| Class | Soluble |
| Log *S* (Ali) | -4.81 |
| Solubility | 9.64e-03 mg/ml ; 1.55e-05 mol/l |
| Class | Moderately soluble |
| Log *S* (SILICOS-IT) | -2.25 |
| Solubility | 3.48e+00 mg/ml ; 5.60e-03 mol/l |
| Class | Soluble |
| **Pharmacokinetics** | |
| GI absorption | Low |
| BBB permeant | No |
| P-gp substrate | Yes |
| CYP1A2 inhibitor | No |
| CYP2C19 inhibitor | No |
| CYP2C9 inhibitor | No |
| CYP2D6 inhibitor | No |
| CYP3A4 inhibitor | No |
| Log $K_p$ (skin permeation) | -9.76 cm/s |
| **Druglikeness** | |
| Lipinski | No; 3 violations: MW>500, NorO>10, NHorOH>5 |
| Ghose | No; 4 violations: MW>480, WLOGP<-0.4, MR>130, #atoms>70 |
| Veber | No; 1 violation: TPSA>140 |
| Egan | No; 1 violation: TPSA>131.6 |
| Muegge | No; 4 violations: MW>600, TPSA>150, H-acc>10, H-don>5 |
| Bioavailability Score | 0.17 |
| **Medicinal Chemistry** | |
| PAINS | 0 alert |
| Brenk | 0 alert |
| Leadlikeness | No; 2 violations: MW>350, Rotors>7 |
| Synthetic accessibility | 6.63 |

Figure S11: Physicochemical and AMDE properties for compound pectolinarin

#### 6.) Used software tools and versions for the virtual screening analyses

**a) Protein structure pre-processing and quality control**

- Schrödinger Maestro software (version 11.8.0.1.2, www.schrodinger.com)
- Verify3D (as implemented in SAVES 5.0, http://servicesn.mbi.ucla.edu/SAVES)
- WHATCHECK (as implemented in SAVES 5.0, http://servicesn.mbi.ucla.edu/SAVES)
- PROCHECK (as implemented in SAVES 5.0, http://servicesn.mbi.ucla.edu/SAVES)

**b) Molecular docking and ligand pre-processing, binding affinity estimation, and ligand similarity screening**

- OpenEye docking software (version: 2019.Nov.2-Ubuntu-18.04-x64), including software tools OMEGA (version: 3.1.2.2) and HYBRID (version 3.4.0.2)
- AutoDock-GPU (version: OpenCL and Cuda accelerated version of AutoDock4.2.6, Linux, x64, autodock_gpu_256wi) and MGLTools package (version 1.5.6, Linux, x64)
- BiosolveIT LeadIT software (version: 2.1.5, Linux, x64), including binding affinity estimation software HYDE (version: 3.2.5)
- BiosolveIT FTrees software (version: 6.2, Linux, x64)

**c) Machine learning based compound screening:**

- R statistical programming software (version: 3.6.0, Linux, x64)
- R software package "rcdk" (version: 3.5.0)
- R software package "rcdklibs" (version: 2.3)
- R software package "iterators" (version: 1.0.12)
- R software package "randomForest" (version: 4.6-14)

**d) Molecular dynamics simulation and molecular visualization software**

- NAMD Molecular Dynamics Software (version: Git-2020-07-30, Linux, x86_64-multicore-CUDA)
- VMD molecular visualization software (version: 1.9.3)
- UCSF Chimera molecular visualization software (version 1.12, build 41623)
- PoseView 2D complex diagram generation and visualization software (version 1.1.2)
- LigandScout software for pharmacophore generation and visualization (version 4.4.5)

### 7.) Step-by-step description of the virtual screening analyses

**1.) Ligand collection, pre-processing and filtering**

a) ZINC database:

Compounds were downloaded in SMILES-format, using the “ZINC-downloader-2D-smi.wget” script derived from the “Tranches” web-page on ZINC (https://zinc.docking.org/tranches/home, version ZINC15, May 2020). The following filtering criteria were applied to all compounds available via the ZINC Tranches web-page: “drug-like”, “purchasable” (minimum purchasability = “Wait OK”) and reactivity = “clean”. This first-step filtering reduced the initial 1,276,766,435 substances to 898,838,573 retained substances.

b) SWEETLEAD database:

All compounds from the SWEETLEAD database (version: 1.0) were downloaded in May 2020 (https://simtk.org/projects/sweetlead).

c) MolPort database:

All compounds from the MolPort library were downloaded in May 2020 (https://www.molport.com/shop/database-download).

d) Ligand pre-processing:

Ligand preprocessing involved adding hydrogens and generating 3D structures, where not already provided in the source databases, while protonation states and partial charges were assigned during the docking stage. Specifically, the compounds were preprocessed using the AutoDock ligand preparation script (prepare_ligand4.py) from the MGLTools package (version 1.5.6, Linux, x64) with default parameters, and conformers were generated using the OpenEye OMEGA software (version: 3.1.2.2) using the classic mode with default parameters.

e) Ligand-based similarity screening / filtering:

Since the number of pre-selected compounds obtained from the ZINC database was too large for comprehensive docking analyses, this compound library was further filtered using a ligand-based similarity screening with the BioSolveIT Ftrees software (version 6.2). The topological and physicochemical similarity of the library compounds to known small-molecule inhibitors for SAR-CoV and SARS-CoV-2 3CLpro reported in the literature was scored with this method, to filter the library down to the compounds most similar to known 3CLpro inhibitors. Specifically, the literature-derived query compounds include the reported SARS-CoV-2 3CLpro inhibitors GC-376, ebselen and baicalein, and the reported SARS-CoV 3CLpro inhibitors amentoflavone, hesperetine, pectolinarin and dieckol. Moreover, to further extend the search space of potential candidate inhibitor compounds, four additional query compounds for the feature tree search were included by adding the top-ranked compounds from the initial AutoDock-GPU screening on the SWEETLEAD database. All compounds from the ZINC library exceeding a minimum similarity threshold of 0.8 in the FTrees screen to these query compounds were retained for the subsequent molecular docking analyses (see script "ftrees_screening.R" on the GitHub web-page https://github.com/eglaab).

**2.) Protein structure selection, pre-processing and quality control**

a) Protein structure selection:

Three publicly available protein crystal structures for 3CLPro from the Protein Data Bank (PDB) were chosen for the molecular docking analyses and binding affinity estimation: 5R8T, 6YB7 and 6LU7. The structures 5R8T and 6YB7 were selected mainly due to their resolution (5R8T: 1.27 Å, 6YB7: 1.25 Å) and quality (R-free value: 0.208 for 5R8T and 0.192 for 6YB7), see additional quality assessments described below), whereas the structure 6LU7 was used additionally as a representation of the holo form of the protein in complex with an inhibitor (resolution: 2.16 Å, R-free value: 0.235), allowing us to compare docking results across different types of structures.

b) Protein structure pre-processing:

The receptor structures were pre-processed using the Schrödinger Maestro software (version 11.8.0.1.2, www.schrodinger.com) by adding hydrogens, generating protonation states, and optimizing hydrogen positions using the ‘Protein Preparation Wizard’ with default settings.

c) Protein structure quality control:

The quality of the original and processed structures was assessed using the software tools Verify3D, WHATCHECK and PROCHECK as implemented in the software SAVES 5.0 (http://servicesn.mbi.ucla.edu/SAVES), confirming the suitability of the structures for docking simulations in terms of common quality control checks [37]. For structure files containing multiple chains, the chain with the highest Verify3D score was chosen for further analysis.

**3.) Virtual screening**

a) Docking using AutoDock-GPU:
The pre-selected compound libraries were docked using AutoDock-GPU (version: OpenCL and Cuda accelerated version of AutoDock4.2.6, Linux, x64, autodock_gpu_256wi) with the parameter 'nrun' for the thoroughness of the search space exploration set to 100, and default parameters otherwise (see script "autodock_screening.R" on the GitHub web-page https://github.com/eglaab).

b) Docking using OpenEye HYBRID:
The pre-selected compound libraries were docked using OpenEye HYBRID with default docking parameters (see script "openeye_hybrid_docking.sh" on the GitHub web-page https://github.com/eglaab).

c) Docking using LeadIT/FlexX + binding affinity estimation with LeadIT/HYBRID:
Only compounds with higher than average docking scores derived from the AutoDock-GPU and OpenEye HYBRID screenings were also docked using a more time-consuming combined docking and binding affinity estimation approach, by applying the software LeadIT/FlexX (version 2019.Nov.2-Ubuntu-18.04-x64) with default docking parameters and a subsequent estimation of the binding affinity for the top 30 docking poses using the LeadIT/HYDE approach (version: 3.2.5).

d) Ranking and selection:
After obtaining the docking scores for each of the three docking approaches, the list of compounds docked with each method was ranked and sorted according to the sum of ranks across the scores for all methods. Only top-ranked compounds achieving consistently high docking scores for the three pre-processed 3CLpro protein structures (PDB: 5R8T, 6YB7 and 6LU7) were used for experimental validation.

e) Alternative screening approaches:
For the MolPort compound library, we tested multiple alternative more extensive screening approaches without prior library filtering: (1) a screening approach relying purely on fast molecular docking approaches (see sections 3 a), b) and c)), (2) a

screening approach relying purely on machine learning (see section 3 f) below), and (3) a combination of molecular docking and ligand-similarity based screening using the software FTrees (following the same approach as for the ZINC database described in section 1 e), but without prior database filtering, and focusing on the most potent available 3CLPro inhibitor, GC-376, as query compound). This extended analysis was limited to the MolPort compounds, because this library was our main resource for commercially available compounds and therefore of particular practical relevance for the experimental studies, and due to its smaller size (~7.7 million compounds), an extension to further screening approaches was still feasible in terms of runtime requirements.

f) Machine learning based screening:
To assess a 3CLpro candidate inhibitor ranking approach using machine learning and extend the selection of compounds for experimental testing, molecular descriptors were computed for the compounds, as implemented in the R software package "rcdk" (version 3.5.0) using SMILES compound representations as input. For the training of the machine learning model, a published training set collection of small molecules containing both previously reported SARS-CoV or SARS-CoV-2 3CLPro inhibitors (486 compounds, used as positive set) and compounds reported not to bind to 3CLPro (700 compounds, used as negative set) was used to compute the moleclular descriptors (Ivanov et al., ACS Omega, 2020). On this training data of descriptors for the positive and negative compound sets, a classification model was trained using the Random Forest algorithm as implemented in the R software package "randomForest" (version 4.6-14) with 250 decision trees and default parameters otherwise. This trained model was used to rank all compounds in the MolPort library, by computing the molecular descriptors with the "rcdk" package and applying the trained model to estimate the probabiliy of belonging to the active inhibitors (see script "ml_screening.R" on the GitHub web-page https://github.com/eglaab).

**4.) Molecular Dynamics simulations**

The molecular dynamics (MD) simulations to analyze the ligand binding stability for the top-ranked compounds obtained from docking simulations in the 3CLpro binding pocket were performed using a GPU-accelerated version of the software NAMD (Git-2020-07-30 for Linux-x86 64-multicore-CUDA). All non-default configuration settings used for the simulations are specified in the file "ligand_binding_simulation.namd" available on the GitHub web-page (https://github.com/eglaab). Briefly, the step size was 2 fs, and the simulation was performed for a duration of 20 ns, the topology and force field parameters for the MD simulations were assigned from the CHARMM36 protein lipid parameter set for the receptor, and from the CHARMM General Force Field (CGenFF) parameter set for the ligands. Video animations of the simulations were created by applying the “MD Movie” function in the software Chimera (version 1.12, build 41623, [51]) to the trajectory data for the protein-ligand complexes derived from NAMD.
